# Boolean dynamic modeling of TNFR1 signaling predicts a nested feedback loop regulating the apoptotic response at single-cell level

**DOI:** 10.1101/2022.07.29.502000

**Authors:** Shubhank Sherekar, Ganesh Viswanathan

## Abstract

Cell-to-cell variability during Tumor Necrosis Factor Receptor 1 (TNFR1) signaling triggered by the pleiotropic cytokine TNFα can lead to pro-survival and apoptotic phenotypic responses at single-cell level. Harnessing the ability to modulate the signal flow responsible for the balance between these two phenotypes and make cells favour apoptosis have been considered in cancer therapies. We show that a 6-node nested feedback loop facilitates the crucial crosstalk regulation modulating the signal flow between these two responses. We identify this by systematically analysing the partial state transition graph (pSTG) underlying a Boolean dynamic model of the TNFR1 signaling network that accounts for signal flow path variability. We demonstrate a novel approach “Boolean Modeling based Prediction of Steady-state probability of Phenotype Reachability (BM-ProSPR)” that enables constructing a reliable pSTG in a computationally efficient manner and predicting accurately the network’s ability to settle into different phenotypes. We deduce that knocking-off Comp1 – IKK* complex tweaks the signal flow paths leading to a 62% increase in the steady state probability of TNFR1 signaling culminating in apoptosis and thereby favours phenotype switching from pro-survival to apoptosis. Priming cancerous cells with inhibitors targeting the interaction involving Comp1 and IKK* prior to TNFα exposure could be a potential therapeutic strategy.

## 1. Introduction

Apoptotic response is a key outcome of Tumor Necrosis Factor Receptor 1 (TNFR1) signaling triggered by TNFα, a pleiotropic cytokine. While normal cells maintain a balance between prosurvival and apoptotic responses due to TNFα,^1,2^ diseased ones such as those of certain cancer tissues favor proliferation by evading apoptosis.^3^ However, counter-intuitively, certain tumors have higher sensitivity to apoptosis than the corresponding normal tissue.^4,5^ Modulation of the delicate balance causing switching of phenotype from pro-survival to apoptotic response can be harnessed for novel therapeutic approaches involving TNFα.^6^ How cells can be primed to exhibit apoptotic outcome due to TNFR1 signaling? Regulating the activity of proteins activated is a priming approach. Identifying these proteins require understanding the signal flow through a cell.

Signal flow through TNFR1 signaling network is orchestrated by several underlying myriad molecular markers.^7,8^ Cell-to-cell variability causing an ensemble-level behavior can influence the signal flow through every cell in a population.^2,9,10^ While TNFR1 signaling has been modeled at both population-averaged and single-cell levels,^11–13^ it is yet unclear what are the relevant markers that together regulate the phenotypic balance at an ensemble-level and how they influence the signal flow. Understanding this cross-talk regulation in a quantitative manner will help identify strategies for signal flow modulation enabling phenotype switching to apoptosis.

Deterministic kinetic modeling of TNFR1 signaling revealed that, at population-averaged level, the pro-survival response, reflected by NFκB activation, is TNFα stimulus strength dependent.^14^ TNFR1 signaling mediated cell death response in hepatocytes is guided by sensitizing the apoptotic mediators^13,15^ such as caspase3 via TRADD complex.^16^ Contrastingly, TNFR1 signaling dictated cell-fate is determined by intracellular protein marker activity indicating that these are signatures of the underlying phenotypic regulation.^17–19^

Kinetic modeling is unsuitable for large networks as it is fraught with unavailability of estimates for most parameters. Boolean dynamic (BD) modeling which assumes every node can be either active (ON) or inactive (OFF) circumvents this limitation.^20,21^ Boolean model of TNFR1 signaling network using synchronous updating scheme showed that feedback loops involving caspase3 may control culmination into either pro-survival or apoptotic response.^22^ Multivalued discrete dynamical simulations, an extension of BD modeling, predicted that hepatocytes and Jurkat-T cells require smac – mimetics for the TNFR1 signaling mediated apoptotic response.^11^ Moreover, pulsatile TNFα stimulation reduces resistance caused by NFκB only in the absence of active IKK.^23^ These models predict reachability to only one phenotypic response from a certain initial condition mimicking the internal state of a cell. However, experimental studies show that individual cells in a population grown under identical conditions and exposed to same stimulus conditions can lead to different phenotypic responses.^24^ The inability of these TNFR1 signaling synchronous updating-based BD models to predict multiple outcomes from an initial state could be attributed to lack of incorporation of cell-to-cell variability.

Use of general asynchronous (GA) or random-order asynchronous (ROA) updating scheme permits incorporation of cell-to-cell variability in the BD model of the network while estimating its ability to settle into different phenotypes.^21,25,26^ GA updating-based BD model of TNFR1 network unraveled that inhibition of AKT/PKB,^27^ RIPK1^28^ and FADD^28^ are crucial in modulating the steady-state probability of attaining long-term pro-survival responses (attractor). Since GA update scheme could introduce spurious self-loops and bias signal flow paths to prefer a certain entity, the probabilities predicted by these models may not reflect the true extent of the network’s ability to settle into the attractors.

The primary goal of this study is to model TNFR1 signaling accounting for cell-to-cell variability in signal flow path using ROA updating strategy, which improves the accuracy of the steady-state probabilities and to distill out the cross-talk signaling machinery regulating the phenotypic responses. While ROA updating-based BD modeling does not lead to spurious self-loops and significantly minimizes biases, it requires construction of the complete state transition graph (STG), a collection of all signal flow paths, which is computationally infeasible even for a reasonably sized network. We develop a systematic approach <u>B</u>oolean <u>M</u>odeling based <u>P</u>rediction <u>o</u>f <u>S</u>teady-state probability of <u>P</u>henotype <u>R</u>eachability (BM-ProSPR) to construct a reliable *partial* STG that can accurately predict the steady-state probabilities of reaching multiple phenotypes. After benchmarking on multiple networks, using the partial STG underlying the BD model of TNFR1 network, we identify the crucial signaling cross-talk regulation. We show that perturbation of signal flow via entities involved in this cross-talk regulation can enable phenotype switching from pro-survival to apoptosis.

## 2. Results

### 2.1: Boolean model of TNFR1 signaling mediated apoptosis

We manually curated a molecular wiring diagram of TNFR1 signaling originating from TNFα and leading to apoptotic response (Fig. 1).^29,30^ Fas Ligand (FasL), a TNF superfamily ligand, too leads to a strong apoptotic response via Fas receptor.^31^ Since Fas mediated signaling shares majority of the death signal pathways in TNFR1 network, we considered apoptotic response originating from FasL as well for benchmarking purposes (Fig. 1). The signaling network consists of *N* = 38 entities and 32 causal interactions connecting them. (A detailed description of the network construction along with a list of entities are in Text S1.1 and Text S1.2, Supplementary Information.) We classify these entities into 9 housekeeping (H), 2 input (I), 25 signaling (S) and 2 output (O) nodes (Table S1, Supplementary Information). While the signal transduction is initiated via one of the input nodes, that is, TNFα or FasL, the housekeeping ones represent those present constitutively. On the other hand, signaling nodes direct and regulate the signal flow through the network to the output nodes Apoptosis and NFκB. The network’s long-time response can be classified into either apoptotic or pro-survival or anti-apoptotic phenotype depending upon the different combinations of the active/inactive form of the output nodes (Table 1).

**Figure 1:**
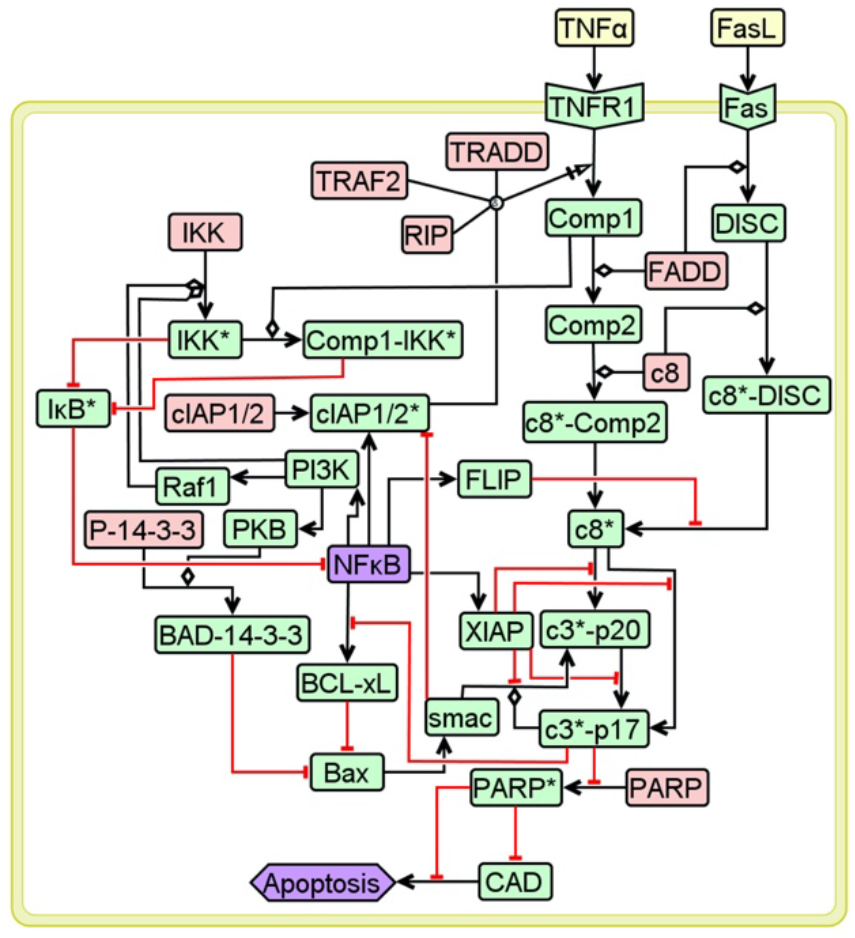
TNFR1 signaling network. Entities in pink, yellow, green and purple, respectively represent housekeeping, input, signaling and output nodes. Black arrows and red hammers capture the activation and inhibitory interactions. Detailed description of the network along with the node-specific Boolean functions are in Text S1, Supplementary Information.

**Table 1:** Phenotypes permitted by the TNFR1 signaling network (Fig. 1) and the conditions that specify them. Values inside a bracket indicate the Boolean value taken by the output nodes Apoptosis and NFκB.

| **Apoptosis**<br>**NFκB** | **Inactive (0)** | **Active (1)** |
| --- | --- | --- |
| **Inactive (0)** | Anti-apoptotic | Apoptotic |
| **Active (1)** | Pro-survival | --- |

We ask a question as to how transient signaling orchestrates the phenotypic response in an ensemble of cells stimulated with TNFα. In order to decode this, we systematically analyze the TNFR1 network dynamics using the Boolean Dynamic (BD) modeling framework, wherein a node *i* can take either active (ON) or inactive (OFF) form quantified respectively by Boolean value *v_i_* = 0 or 1. While *v_i_* for housekeeping nodes are fixed as 1, that for an input entity is set to 1 depending on the stimulation condition considered. Using appropriate Boolean logic to mimic the causal interaction between the entities, we define Boolean functions *f_i_* corresponding to a signaling/output node *i*.^20^ For example, as entity c8* is activated by c8* – DISC in the absence of FLIP or by c8* – Comp2, the corresponding Boolean function is specified by

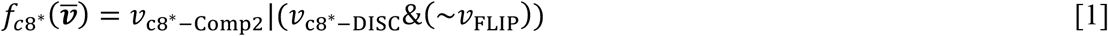

where, set 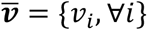 captures the instantaneous *state* of the network mimicking that of a cell. Note that in Eq. (1), &, | and ~, respectively represent AND, OR and NOT logic. While 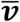 is in general an unordered set, for ease of representation and analysis, we place them in a specific order: 9H, 2I,25S, 2O (Table S1, Supplementary Information). Logic adopted for the interactions are incorporated in the 27 node-specific Boolean functions, as detailed in Text S1.2, Supplementary Information.

Starting from a randomly chosen initial state 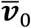, we perform Boolean dynamics of the network using asynchronous updating of the Boolean functions till an absorbing state is reached. (Note that asynchronous updating enables capturing behavior of an ensemble of cells, wherein individual cells initially at a certain state can choose its fate resulting in a cell-specific phenotype.^25^) Such an absorbing state can be either a fixed point consisting of a single state or a cyclic attractor, which meanders over a set of states.^20^ In this study, we only consider the network reaching fixed point attractors (FPs). Values of *v*_Apoptosis_ and *v*_NFκB_ that specify the three attractors are in Table 1. A set of states traversed between 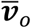 and a FP specifies the corresponding signal flow path to an attractor. A directed transition between two consecutive states in this path is called a one-step state-transition. An input condition- and Boolean function-specific state transition graph (STG), whose state-space 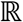 consists of 2*^N^* states, is a collection of all directed signal flow paths to various attractors from any 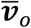 permitted.

We hypothesize that a cell-specific response is caused by the cell-to-cell variability in the signal flow path taken by it to reach a certain phenotype upon stimulation. Random order asynchronous (ROA) method of updating offers mimicking stochastic behavior due to interdependency of the nodes at all update steps and thereby permits extensive intricate signal paths to reach an attractor from an initial state. The variability due to the interdependency of the nodes is achieved by choosing a random sequence in which the Boolean functions are evaluated at every one-step state transition. As a consequence, FP reached by following a signal flow path involving several state transitions strongly hinges on the permutations that specify this random sequence. We therefore adopt ROA update strategy for BD simulations (Methods).

We first reduce the dimensionality of the network by performing a partial-Logical steady state analysis (pLSSA)^32^ (Methods; Text S1.3, Supplementary Information). pLSSA fixes values of certain nodes due to the underlying input condition-specific Boolean functions dictating a partial steady-state and reduces the number of dynamically varying signaling entities. For e.g., *v*_TNFα_ = 1 implies *v*_TNFR1_ = *v*_Comp1_ = 1 based on their Boolean functions (Table S2, Supplementary Information). Under TNFα stimulation conditions, pLSSA led to fixing the Boolean values of 10 signaling nodes, *viz*., TNFR1, Comp1, Comp2, c8* – Comp2, Fas, DISC, c8* – DISC, c8* – cIAP1/2*, c3* – p20, values for which are in Table S3, Supplementary Information and thereby resulting in 17 dynamically varying signaling/output entities. Note that the value of these “fixed nodes” remain unchanged even if that of the remaining entities vary dynamically. The network (Fig. 1) stimulated with TNFα can lead to *only* two FPs, *viz*., pro-survival FP 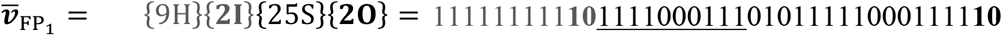 and apoptotic FP 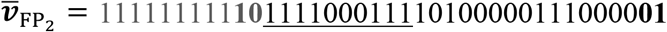, where entities corresponding to those underlined digits are pLSSA-fixed (Methods; Text S2, Supplementary Information). FPs for stimulation with TNFα, FasL and for no stimulation are in Text S2, Supplementary Information. For the sake of brevity, the first 9 digits (housekeeping, fixed at 1) in 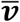 will henceforth be collectively represented as “H”. Starting from an initial state 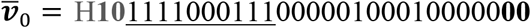, chosen uniformly randomly, BD simulations led to either 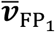 or 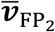 depending upon the signal flow paths dictated by the chosen permutations (Fig. 2A). There could be several signal flow paths leading to a certain FP from an initial state. Since stochasticity in phenotypic response originates from the cell-to-cell variability in signal flow paths, knowledge of these will enable computing the absorption probability of the network to reach a FP starting from a state 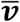. Stimulation condition-specific steady-state probability 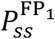 of the network to settle into FP_1_ is given by

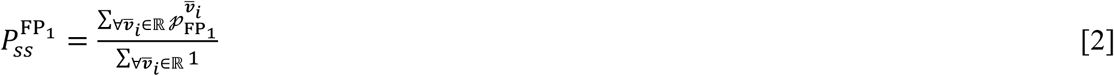

where, 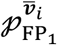 is the absorption probability with which a state 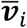 settles into FP_1_. Computing these absorption probabilities requires reliable estimation of the state transition matrix (*M*) quantifying the corresponding STG. An element *M_ij_*, the transition probability associated with the one-step transition between states *i* and *j*, is given by

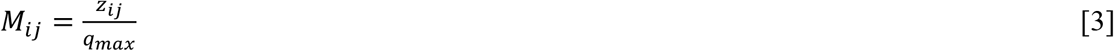

where, *z_ij_* is the number of permutations causing one-step transition between *i* and *j*, and *q_max_* = ∑_∀*,j*_ *z_ij_* is the maximum possible permutations. Computing *M_ij,_* ∀*i, j* involves repeated evaluation of 17 one-step state transition causing Boolean functions for all possible permutations. We therefore ask a question what is the absorption probability of the TNFα stimulated network (Fig. 1) to reach the two FPs from all possible initial states, each of which, chosen uniformly randomly, mimics the internal state of a cell in an ensemble.

**Figure 2:**
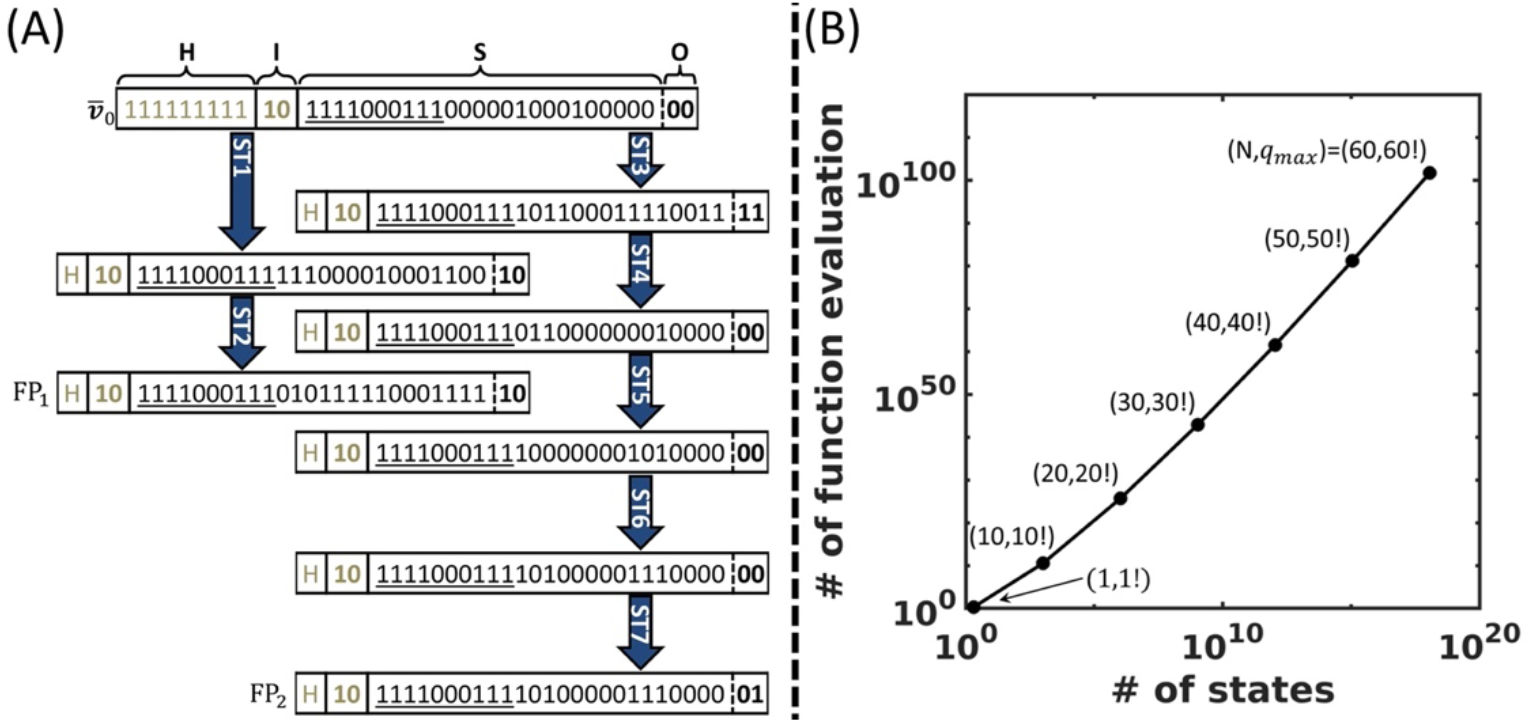
**(A) Absorption of the initial state 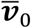 into two fixed points via distinct signal flow paths**. The order in which the Boolean values of the entities are placed in a state is housekeeping (H), input (I), signaling (S) and output (O) nodes. Blue arrow indicates one-step transition with a certain permutation using Random Order Asynchronous (ROA) update scheme. Permutation used in the Boolean simulations of the one-step transitions are in Text S2.2, Supplementary Information. **(B) Effect of the number of entities on the required number of function evaluations for the construction of the state transition graph.**The two numbers within the brackets capture the number of entities and the corresponding number of permutations.

Quantifying *M* for TNFα stimulated TNFR1 network demands 2^*N* = 17^ × 17!× 17 (no. of states × no. of permutations/state × no. of Boolean function evaluations/permutation) Boolean function evaluations. Moreover, every function could in turn have several Boolean operations embedded in it depending on the network wiring. Therefore, for any general network, the number of Boolean function evaluations required for estimating *M* increases exponentially with *N* (Fig. 2B). Thus, computing *M* is prohibitively expensive, in spite of reducing the network dimensionality and fixing the housekeeping and input nodes. Therefore, estimation of the state transition matrix (*M_c_*) of the complete STG using ROA update strategy is *infeasible* even for a reasonably sized large network. This challenge can be circumvented if the redundancy in the signal flow paths to a FP can be minimized while preserving the transition probabilities in *M*. This redundancy stems from the fact that many chains of permutations could lead to identical signal flow paths, that is, consisting of the same set of states in the STG. One possible way to reduce the redundancy and thereby the underlying computations is to consider only a certain randomly chosen fraction of maximum possible permutations that predicts a reduced *M* ≈ *M_c_*. In the next section, we develop a *novel* systematic algorithm to identify the fraction of *q_max_* that can reliably quantitate the state transition matrix *M* when ROA update strategy is used. In the subsequent section, we implement this algorithm on the network in Fig. 1 to unravel the signaling transients leading to a phenotypic response due to TNFα.

### 2.2 Boolean Modeling based Prediction of Steady-state probability of Phenotype Reachability (BM-ProSPR)

The goal of BM-ProSPR algorithm is to find 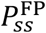 of reaching different FFs by estimating a reliable state transition matrix *M* using *q_l_*, a fraction of the maximum possible permutations *q_max_*. For demonstration purposes, as a motivating example, we consider a small 6-node apoptosis network of T-cell Large Granular Lymphocyte (T-LGL) cells (Fig. 3A).^33^ This network consists of S1P, FLIP, FAS, CERAMIDE, DISC, and Apoptosis –order of appearance of the entities in 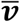. Functions governing the BD of this network (Fig 3A) are in Fig. 3B. T-LGL network permits only two FPs, *viz*., 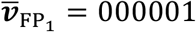 and 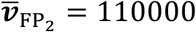, respectively representing apoptotic and anti-apoptotic responses.^33^ Construction of the complete STG being feasible for this small network makes it suitable for benchmarking purposes.

**Figure 3:**
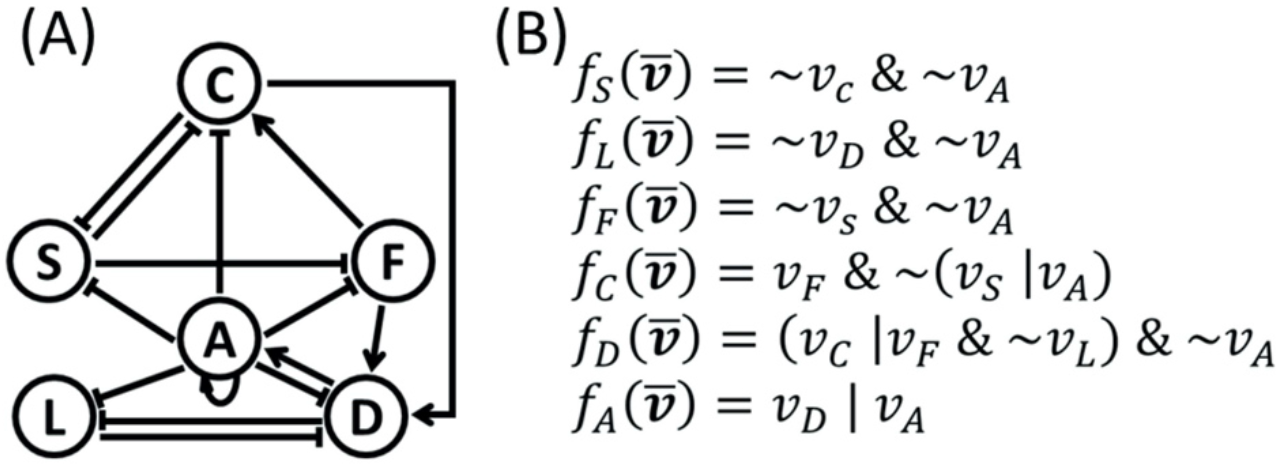
**(A) 6-node T-cell Large Granular Lymphocyte (T-LGL) apoptosis network**. Nodes A, C, D, F, L, and S, respectively represent Apoptosis, Ceramide, DISC, Fas, FLIP, and S1P. Arrows and hammers, respectively capture activation and inhibition interactions. **(B) Boolean functions corresponding to the nodes in (A).** &, |, and ~, respectively correspond to *AND, OR, and NOT* logics.

#### BM-ProSPR Algorithm

Flow chart capturing the systematic approach involved in BM-ProSPR for computing 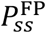 is in Fig. 4. As a first step (Step 1, Fig. 4), we specify the network of interest consisting of N nodes, edges between the entities and node-specific Boolean functions *f*. In the next step (Step 2, Fig. 4), starting with a null STG (*S*^0^) consisting of 2^6^ = 64 isolated states and assuming an initial seed number of permutations *q* = *q*_0_ ≥ 1, we construct a partial STG *S*^*q*_0_^. The steps involved in this construction is in Table 2. After constructing *S*^*q*_0_^, the associated state transition matrix *M*^*q*_0_^, where 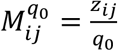 is computed. We show in Fig. 5A snapshots of partial STGs of the 6-node network (Fig. 3) corresponding to 0 (null), 1, and 2 permutations. While *S*^0^ (Fig. 5A-i) consists of only isolated states marked with randomly chosen IDs, *S*^1^ contains 64 one-step transitions (Fig. 5A-ii). These 64 directed links are either self-links from two FP states (state IDs 2 and 49, green circles), or transition into one of the FP states (for e.g., 4 to 2, 33 to 49) or transition between non-FP states (for e.g., 3 to 33). Fig. 5A-ii also shows signal flow paths (for e.g., 5 → 12 → 2, and 3 → 33 → 49) emanating from a state, traversing through other states and culminating in either of the attractors. Upon introduction of a second permutation, 64 new one-step transitions between as many pairs of states are added to *S*^1^ leading to *S*^2^ (Fig. 5A-iii). Out of these 64 pairs in *S*^2^, 24 of them (blue arrows) did not have a link in *S*^1^, while 40 of them (red arrows) accounted for the second one-step transitions between them. The other 24 links (black arrow) in *S*^2^ correspond to those pairs between which a transition was *not* caused by the second permutation and therefore, are carried forward from *S*^1^.

**Figure 4:**
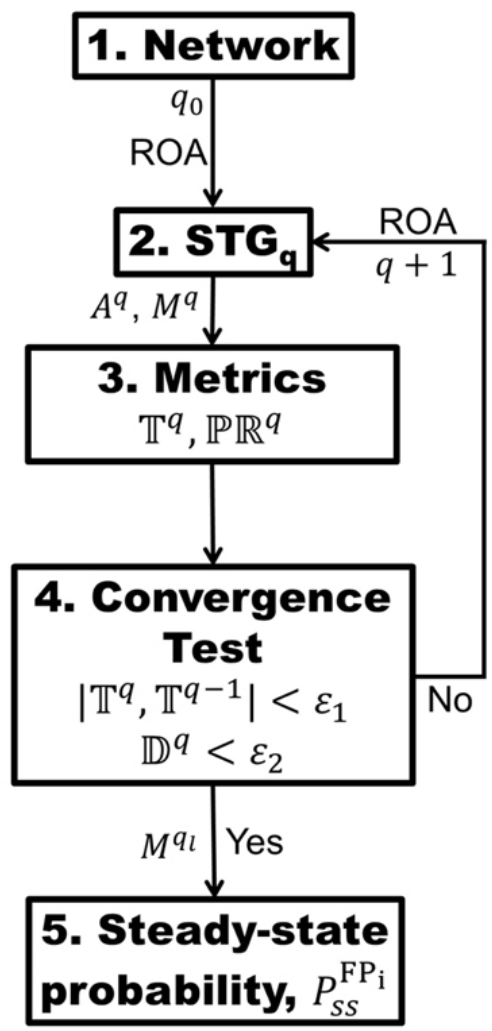
Flow chart elucidating steps involved in BM-ProSPR. While 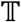 and 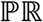, respectively represent Temporality and PageRank measures, 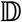 reflects the discordant PageRank fraction across successive permutations. *A^q^* and *M^q^*, respectively are the adjacency matrix and partial STG at the *q^th^* permutation. *ϵ*_1_ and *ϵ*_2_ are the error thresholds. 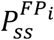 is the steady-state probability estimated using *M^q_l_^* at the minimum required number of permutations *q_l_*. ROA update scheme was used in the BD simulations.

**Figure 5:**
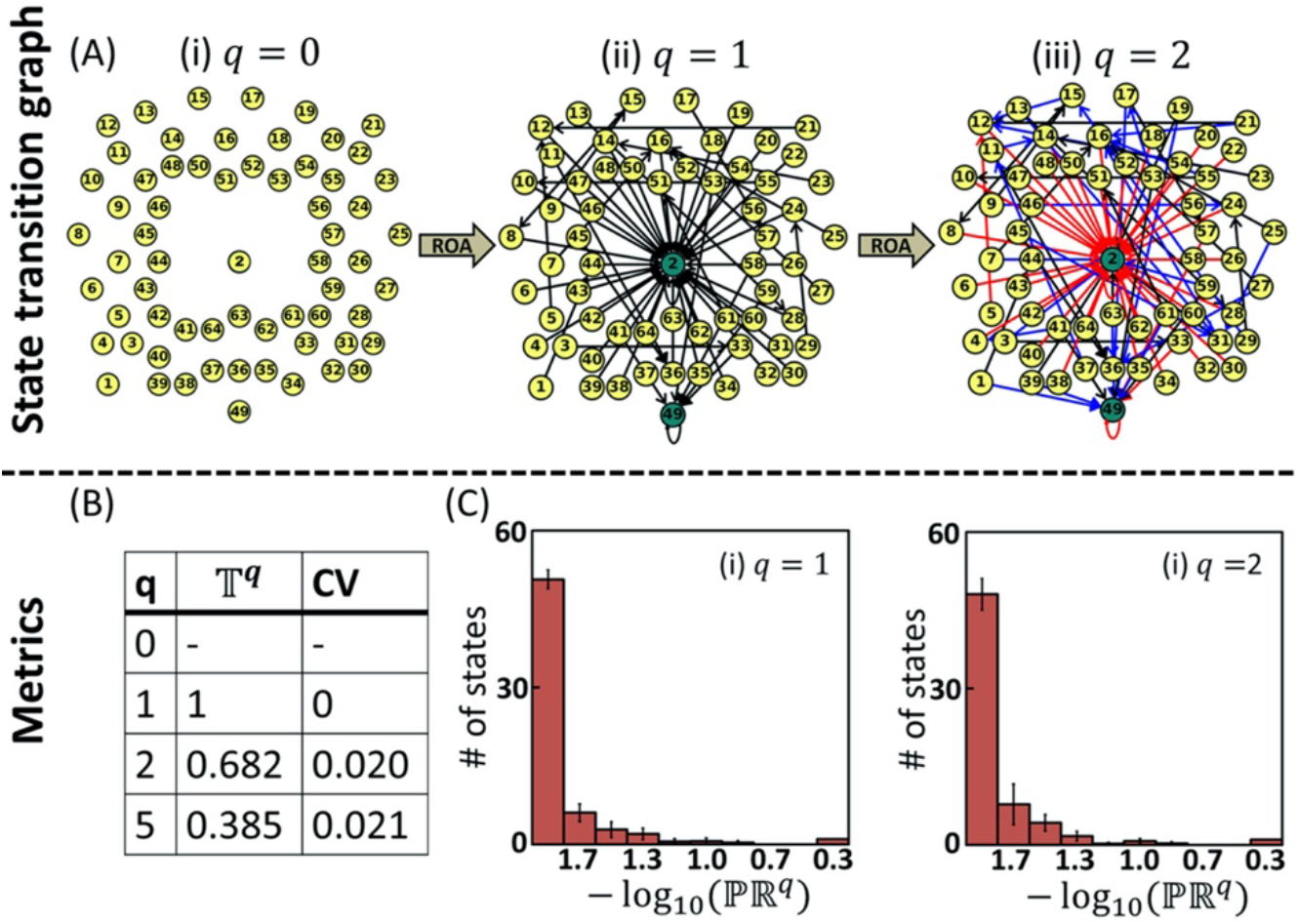
Permutation guided evolution of the state transition graph (STG) underlying the Boolean model of 6-node T-LGL apoptosis network. (A) STG for 0 (null), 1 and 2 permutations. Note for every permutation, 64 new one-step transitions are added. For every one-step transition, BD simulations were performed using a permutation chosen uniformly randomly from 6! (= 720) sequences. Green and yellow circles capture the fixed points and other states, respectively. Black arrows in STG for *q* = 2 are those one-step transitions between a pair of states carried forward, as is, from that constructed for the previous permutation (*q* = 1. While blue arrows represent one-step state transition obtained between a pair of states due to permutation *q* = 2 but not found in the STG for *q* = 1, red arrows capture two one-step transitions, one each obtained between same pair of states for *q* = 1 and *q* = 2. (B) Temporality measure 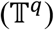 of the STG constructed for a few permutations. The values reported for are 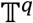 averaged quantities over 50 random constructions of the STG for every *q*. The CV reflects the coefficient of variation of 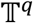 over these random constructions. (C) Histogram of PageRank 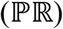 of the states in STGs for *q* = 1 and 2. Note that the mean frequency and the corresponding standard deviation for every bin were estimated using 50 STG reconstructions.

**Table 2:** Steps involved in construction of partial state transition graph (*S^q^*) at *q^th^* permutation.

| Steps involved in constructing $S^q$ |
| --- |
| a) Choose a state $\bar{v}_i \in \mathbb{R}$. |
| b) Choose a permutation from $q_{max} = 6! = 720$. |
| c) Starting from $\bar{v}_i$, for the chosen permutation, identify the state $\bar{v}_e$ reached after one BD simulation with ROA updating strategy (Methods). |
| d) Update $S^q$ with the one-step transitions between states $\bar{v}_i$ and $\bar{v}_e$. |
| e) Repeat steps (b) to (d) for the remaining $q - 1$ times while ensuring no redundancy in the permutation chosen in step (b) for a specific state $\bar{v}_i$ |
| f) Repeat steps (b) to (e) for all $\bar{v} \in \mathbb{R}$. |

Finding 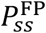 hinges on identifying the minimum number of permutations *q_l_* < *q_max_* such that *M^q_l_^* ≈ *M^q_max_^*, the state transition matrix of *S^q_max_^*. This implies that when *q* = *q_l_* the corresponding partial STG *S^q^* must have sufficiently evolved to contain enough number of directed links such that *S^q^* mimics *S^q_max_^*. We assess the extent of evolution of *S^q^* by monitoring (i) the connectedness between a pair of states and (ii) the number of permutations causing one-step transitions between a pair of states (Step 3, Fig. 4). First, we consider connectedness by tracking the fractional increment in finding at least one directed link between a pair of states resulting from the update due to *q^th^* permutation. We quantitate this fractional increment across permutation-driven STG snapshots using a temporality measure^34–36^

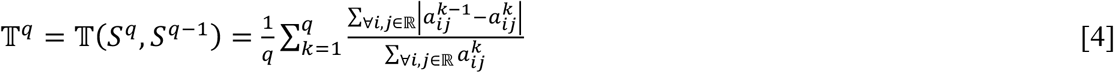

where, 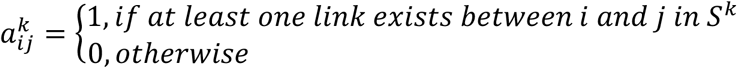 and |.| represents *mod*. 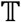 is normalized and 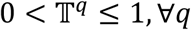 with 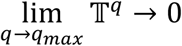, indicating that the STG does not change any further. 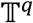 corresponding to the partial STGs in Fig 5A shows a decreasing trend with increasing permutations (Fig. 5B). Next, we quantify the number of permutations leading to a transition between a pair of states by assessing the PageRank 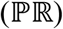^37–39^ of all 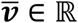 (Methods). 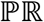 of all states in the null STG is 0. A comparison of the histogram of 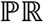 of all 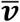 in *S*^1^ and *S*^2^ suggests that addition of 64 new one-step transitions causes a change in the topology (Fig. 5C-ii and 5C-iii), but does not reveal how it led to a change, if any in the order of ranking of the states.

We next introduce conditions on the measures 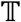 and 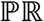 for identifying *q_l_* required to ensure sufficiency in the extent of evolution of the partial STG (Step 4, Fig. 4). *q_l_* required for this sufficiency and thereby, for *M^q_l_^* ≈ *M^q_max_^* is that *q* which satisfies the conditions

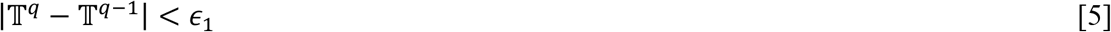

and

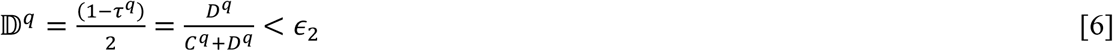

where, 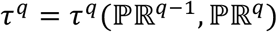 is the Kendall-Tau rank correlation,^40,41^ and *D^q^* and *C^q^*, respectively capture the number of pairs having dissimilar and similar rank-order in 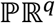 with respect to those in 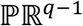 (Methods). Therefore, *C^q^* + *D^q^* = 2^*N* – 1^(2^*N*^ – 1) (Text S3, Supplementary Information). Note that 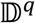(Eq. 6) specifies the fraction of pair of states having discordant PageRank across successive permutations. Thus, 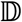 offers a rational comparison of 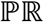 order achieved in successive permutations. We set the error thresholds *ϵ*_1_ and *ϵ*_2_ in Eqs 5 and 6, respectively to ~10^-4^. Conditions in Eqs 5 and 6, respectively ensure that the fractional increment of directed links in and variation in the topology of the STG beyond *q_l_* are insignificant. For a certain *q* if Eqs 5 and 6 are not satisfied, we introduce additional permutation(s) and repeat Steps 2-4 (Fig. 4) until convergence is achieved. After identifying *q_l_*, in Step 5 (Fig. 4), we estimate the steady-state probability 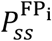 (Eq. 2, Methods).

#### Benchmarkins of BM-ProSPR

We next implemented the entire BM-ProSPR (Fig. 4) on the 6-node T-LGL network by assuming *q*_0_ = 1 (seed permutations) and identified *q_l_* needed to reliably estimate 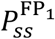 and 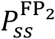. We tested and benchmarked BM-ProSPR by implementing the algorithm for all 720 (= *q_max_*) permutations, rather than terminating at the minimum required permutations *q_l_*. Since the algorithm employs permutations chosen uniformly randomly, we implemented the BM-ProSPR on the T-LGL network 50 times, and estimated 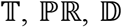 and *q_l_* for each of these. Dependence of the 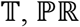 and 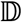 (averaged over 50 STG reconstructions) on the permutations are in Fig. 6A, 6B, and 6C, respectively. We contrast the variation of 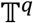 (Fig. 6A, blue) with the distance measure (Fig. 6A, red) given by

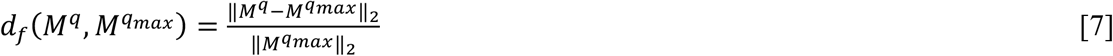

where, ∥.∥_2_ is the Fröbenius norm. *d_f_*(*M^q^, M^q_max_^*) quantitates the extent to which approximates, decreases with increasing *q* and is zero for *q* = *q_max_* (Fig. 6A, red). This comparison clearly shows that 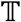 (Eq. 4) is indeed a better indicator of the evolution of the partial STG for a wide range of permutations and accurately reflects the extent to which may have evolved to mimic the complete STG *S^q_max_^* such that *M^q^* ≈ *M^q_max_^*. Moreover, temporality measure (Eq. 4) offers a unique advantage over Fröbenius norm-based distance as 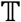 is amenable for large networks for which complete STG and thereby *M^q_max_^* are unavailable.

**Figure 6:**
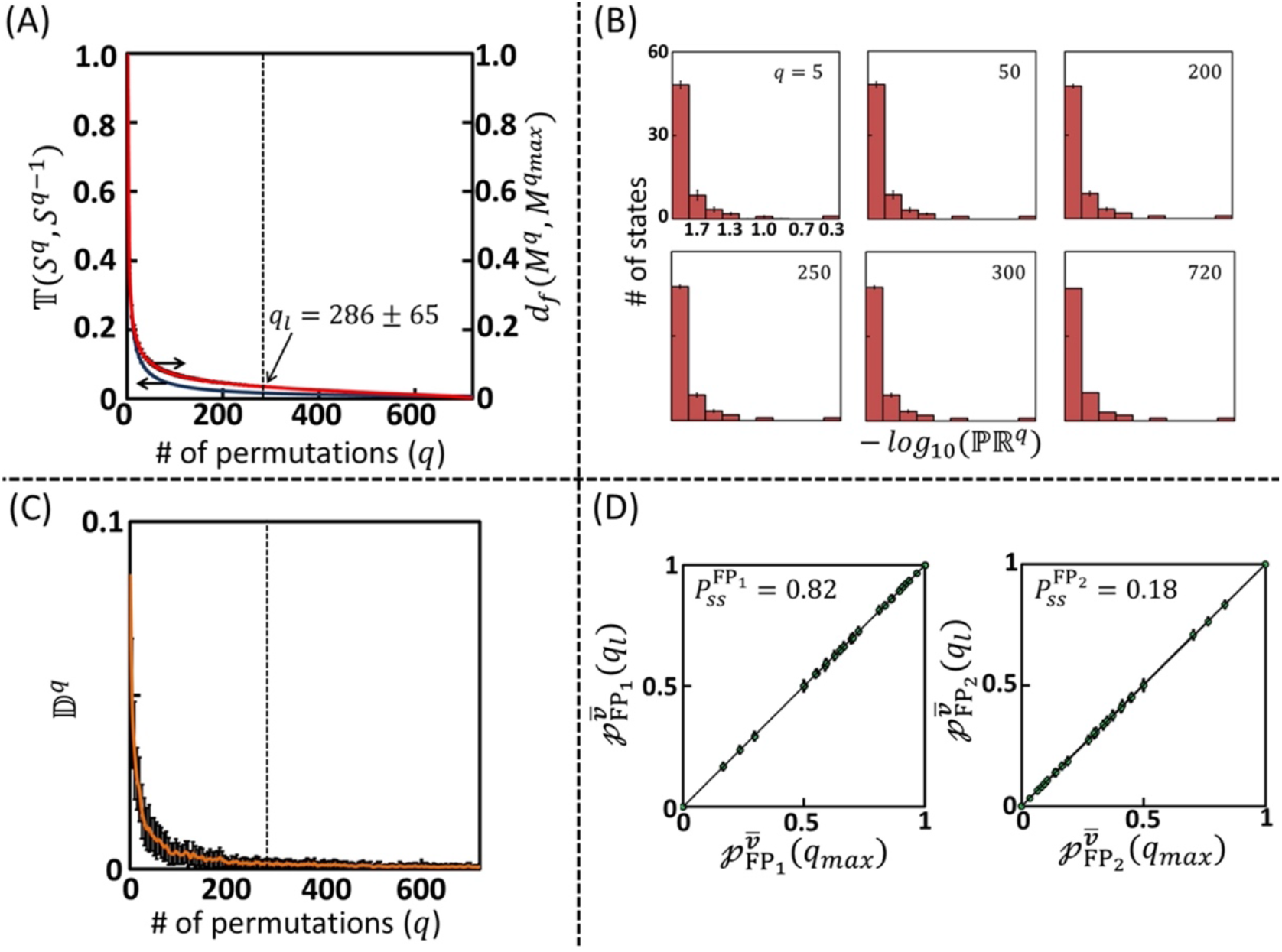
(A) Comparison of the dependence of the Temporality measure 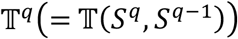 (blue) and the Fröbenius distance *d*(*M^q^, M^q_max_^*) (red) on the number of permutations *q*. *M^q^, M^q_max_^* and are the state transition matrices corresponding to the partial STG at *q^th^* permutation and the complete STG at *q_max_* = 720. *q_l_* is the number of permutations required to construct reliable partial STG. (B) Dependence of the distribution of PageRank 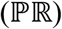 of states on permutations. (C) Dependence of the discordant PageRank fraction across successive STG constructions with increasing number of permutations. (D) A comparison of the absorption probabilities computed using the partial STG at *q_l_* with those specified by complete STG at *q_max_* = 720 to reach the two attractors (FP_1_ and FP_2_) from 64 states. The fractions specified along with the comparison specify the steady-state probabilities of the T-LGL apoptosis network to reach the two attractors. Deviation from the ensemble average of the measures and absorption probabilities obtained over 50 STG reconstructions are presented as error bars.

A comparison of the histograms for different permutations suggests that the 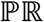 distribution saturates after *q* = 200 indicating that *M*^*q*≥200^ ≈ *M^q_max_^* (Fig. 6B). This is substantiated by the sharp decrease in 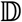 with increasing *q* and insignificant variation for *q* ≥ 200 (Fig. 6C). The conditions in Eqs (5) and (6) require that *q_l_* = 286 ± 65. Note that the relatively large coefficient of variance (CV) across instances of ~23% is due to the tiny statespace of the underlying STG. (We show in the future sections that for large networks, the CV is indeed very small.) Assuming that *S*^286^ ≈ *S^q_max_^*, we estimated the steady-state probability 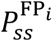 (Eq. 2) to reach FP_1_ and FP_2_ as 0.82 and 0.18, respectively.

For an accurate prediction 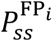, *i* = 1,2, absorption probabilities to reach FP_i_ computed from *M*^286^ must converge to those estimated using *M^q_max_^*. A state 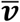 having 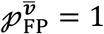 belongs exclusively to the basin of attraction of 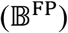. Out of 64 states in 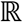, 36 and 3 of them respectively belong exclusively to the basins of attraction of FP_1_ (000001) and FP_2_ (110000) (Table S4, Supplementary Information). The remaining 25 states can reach either of the FPs, each with a finite absorption probability. Absorption probabilities computed from *M^q^* estimated by BM-ProSPR and *M^q_max_^* for both FPs are contrasted in Fig. 6D. This clearly proves that BM-ProSPR indeed predicts the absorption probabilities and therefore, the steady-state probabilities of the 6-node network’s ability to settle into the two fixed points.

BM-ProSPR implemented on an 8-node network regulating the spinal cord ventrilization^42^ permitting five phenotypes showed that the algorithm is effective even when multiple outcomes, a common feature in biological systems, are present (Fig. S4B and Text S5, Supplementary Information). In summary, BM-ProSPR algorithm accurately predicts the steady-state probability of a network settling into different attractors by constructing a partial STG whose state transition matrix mimics that of complete STG. Moreover, temporality measure quantifies the extent of evolution of the STG better even in the absence of a complete one, which is typically the case for large biological networks.

### 2.3 Phenotype switching from pro-survival to apoptosis

We next implement BM-ProSPR on the TNFR1 signaling network (Fig. 1) to decode TNFα mediated apoptotic and pro-survival responses, and identifying targets for phenotype switching. After fixing housekeeping nodes (Table S1, Supplementary Information), we specify *v*_TNFα_ = 1 and *v*_FasL_ = 0. Note that 10 signaling nodes attain pLSS resulting in 17 dynamically varying entities (Section 2.1). BM-ProSPR predicted that 216 (out of 17! = 3.55 × 10^14^) permutations is sufficient to reliably estimate the steady-state probability of reaching the pro-survival 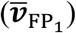 and apoptotic 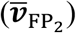 attractors (Fig. 2A). (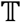 and 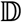 as a function of *q*, and the 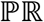 distribution for STG constructed using 216 random permutations are in Fig. S5, Supplementary Information.) TNFα stimulation led to a 20% increase in the steadystate probability (over basal) in the network culminating into a pro-survival phenotype (Fig. 7A). On the contrary, apoptosis phenotype witnessed a significant (7 fold) increase indicating that TNFα signaling drives cell-death response. Note that the anti-apoptotic phenotype (FP_3_), one of the basal responses having a 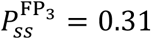, is absent when for TNFα stimulation case.

**Figure 7:**
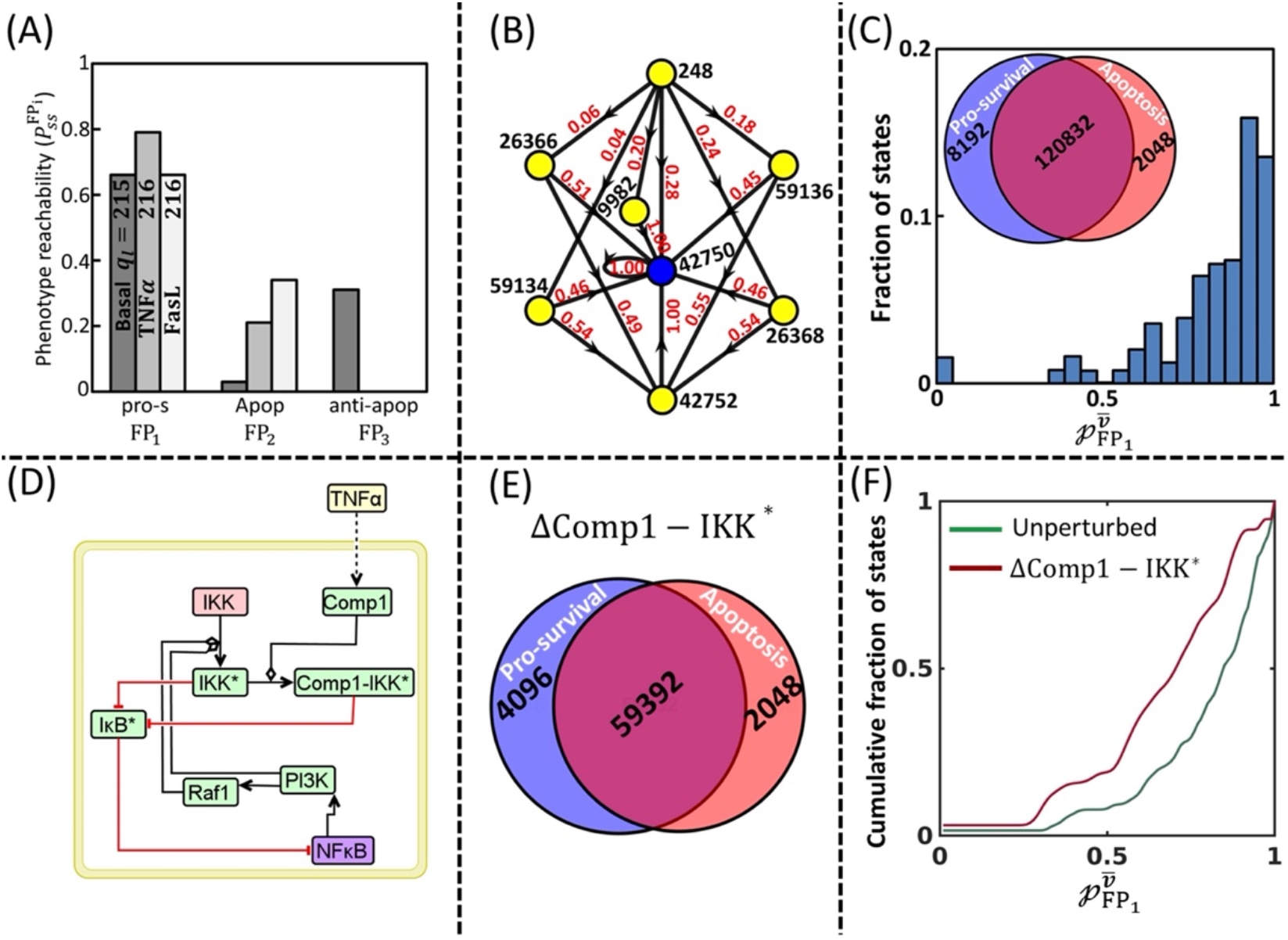
Phenotype switching during TNFR1 signaling. (A) Steady-state probabilities of to reach pro-survival and apoptosis fixed points due to TNFαstimulation. These are juxtaposed with those obtained under no stimulation conditions (basal) and FasL stimulation (positive control). The minimum number of permutations (*q_l_*) required for reliable estimation of the steady-state probabilities to reach the FPs for the case of no stimulation, TNFα, and FasL are 215 ± 0, 216 ± 0, and 216 ± 0, respectively (Text S6, Supplementary Information). Note that under no stimulation conditions, the TNFR1 signaling network (Fig. 1) permitted three FPs, *viz*., pro-survival, apoptotic and anti-apoptotic. Boolean values of the states taken by these FPs are specified in Text S2.1, Supplementary Information. (B) An illustration of the signal flow paths to reach from a randomly chosen initial state (ID-248) to a pro-survival fixed point (FP_1_; ID-42750, blue) for the case of TNFα stimulation. The transition probability associated with each of the one-step transitions are specified next to the corresponding directed link. (C) Distribution of the absorption probabilities of the states to reach pro-survival FP for TNFα stimulation. Inset: Venn diagram showing the number of the states belonging exclusively to the two basins of attraction and those shared between them. (D) Subnetwork showing the nested loop formed by the 6 key nodes regulating the signaling cross-talk in TNFR1 network. (E) Venn diagram showing the number of the states belonging exclusively to the two basins of attraction and those shared between them when Comp1 – IKK* is knocked-off from the TNFR1 signaling network. (F) Comparison of the cumulative distribution of the absorption probability to reach pro-survival phenotype obtained when the TNFR1 signaling network with and without Comp1 – IKK* is stimulated with TNFα.

Under no stimulation conditions, when NFκB is not activated, positive feedback loop involving c3* – p20, c3* – p17 and smac permits c3* – p17 being either active (1) or inactive (0) leading respectively to apoptotic or ant-apoptotic phenotypes. However, when stimulated with TNFα, c3* – p17 can only take an active form in the absence of NFκB. An independent activation of c3* – p20 by FasL (positive control) suggests that anti-apoptotic phenotype is not found when the positive feedback loop is activated under stimulatory conditions (Fig. 7A; Text S6, Supplementary Information).

The orchestration of the TNFα mediated signal transduction leading to phenotypic responses (FPs) is dictated by the absorption probability of different states to reach the corresponding attractor. For example, reaching pro- survival phenotype (ID 42750, FP_1_) from state ID 248 can occur via 10 signal flow paths, each of which has different absorbing capability. While the probability of state 248 to absorb into FP_1_ via path 248 → 59134 → 42752 → 42750 is 0.021(= 0.04 × 0.54 × 1.00), its absorption probability 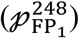 given by the sum of individual path-specific probabilities is 1. We computed 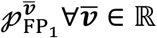 (Methods). Distribution of these 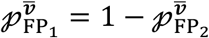 is in Fig. 7C. While 8192 states having 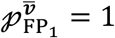 belonged only to pro-survival basin of attraction 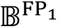, 2048 states with 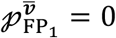 cannot reach FP_1_ (Fig. 7C, inset). The remaining 1,20,832 states belong to both 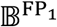 and 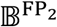 (Fig. 7C, inset). Since each state represents an individual unstimulated cell’s internal condition, the histogram of absorption probability reflects the TNFα mediated phenotypic response of an ensemble of cells. This leads to a question as to what causes a cell having a certain initial state to reach a FP with a certain absorption probability. Deciphering this causality can aid in identifying the network perturbations for modulating a cell’s reachability to a phenotype, particularly that of apoptosis, as it can enable phenotype switching from pro-survival to cell death.

We next systematically analyze the signal flow paths in the STG to unravel the causal effects pertaining to the pro-survival and apoptotic phenotypes. We enumerated the occurrence of nodes with a certain Boolean value in the states that exclusively belonged to the basin of attraction of a FP. For the apoptosis attractor (FP_2_), Boolean value of 6 nodes, *viz*., Comp1 – IKK*, IκB*, IKK*, PI3K, NFκB and Raf1 (Table 3A), which are locked in nested loops (Fig. 7D), were the same in all 2048 states having 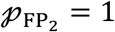. In order for the signal to flow towards apoptosis phenotype, it is necessary that NFκB is arrested as it inhibits cell-death via multiple pathways such as via XIAP, FLIP. Absence of active PI3K and Raf1 results in lack of activation of IKK*, and as a direct consequence, the inhibitory action on IκB* is absent.^43–45^ The active form of IκB is thus responsible for preventing NFκB activation and thereby leading to arrest of signal flow towards pro-survival phenotype (Fig. 1).^46,47^ Out of the 63 other combinations of the value of these 6 key nodes, four of them point to states that belong exclusively in the 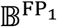(Table 3B). (Note that the remaining 59 combinations have 0 < *p*_FP_1__ = (1 – *p*_FP_2__)< 1.) In all four cases, IκB* takes an inactive form and thereby permitting activation of NFκB leading to pro-survival response. Since inhibitory action on IκB* is either via IKK* or Comp1 – IKK*, IKK* taking a Boolean value of 1 (active) is sufficient to maintain a pro-survival response (Fig. 1). States belonging exclusively to 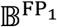 can therefore have either active or inactive Comp1 – IKK*. Thus, these 6 nodes locked in the nested loop regulate the cross-talk signaling between the pro-survival and apoptotic responses.

**Table 3:** Combination of Boolean values taken by the 6 key nodes in the states that belong exclusively to either (A) apoptosis or (B) pro-survival attractors.

| Comp1<br>– IKK* | IκB* | IKK* | PI3K | NFκB | Raf1 |
| --- | --- | --- | --- | --- | --- |
| **(A) Apoptosis attractor (FP<sub>2</sub>)** |  |  |  |  |  |
| 0 | 1 | 0 | 0 | 0 | 0 |
| **(B) Pro-survival attractor (FP<sub>1</sub>)** |  |  |  |  |  |
| 0 | 0 | 1 | 1 | 1 | 0 |
| 0 | 0 | 1 | 1 | 1 | 1 |
| 1 | 0 | 1 | 1 | 1 | 0 |
| 1 | 0 | 1 | 1 | 1 | 1 |

Switching of phenotype between the FPs can be enabled by either (i) shifting an initial state belonging exclusively to the 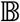 of one FP into another or (ii) increasing the absorption probability of an initial state to a desired FP. Switching from pro-survival to apoptosis via the first option can be achieved by fixing the Boolean values of the 6 cross-talk regulating nodes to that in Table 3A. However, only 6% of the state-space belong uniquely to 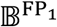, implying that only a small fraction of the cells in an ensemble can be switched. On the contrary, 92.2% of the state-space, that is, majority of cells in a population, can take either of the FPs with a finite probability. Therefore, we consider the case of enabling switching by increasing 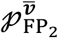. Eliciting a switch in the phenotypic response will require introduction of network perturbations, which we identify by analyzing the signal flow paths taken by the 1,20,832 states that belong to both basins of attraction.

Out of 1,22,880 (= 1,20,832+2048) states belong to 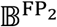, 37,761 states absorb into apoptotic phenotype via a single one-step transition (direct link). Of these, only 8 (including FP_2_) are exclusive to 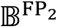. The remaining 85,119, having a finite probability 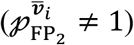 of reaching FP_2_ must culminate into apoptosis attractor via at least one of these 37,761. This suggests that the signal flow path for a state absorbing into apoptotic attractor must either (a) have *at least* one intermediate state having the values of the cross-talk signaling nodes specified by Table 3A or (b) have the *last* one-step transition leading to apoptosis attractor causing the cross-talk signaling nodes taking the values specified by Table 3A. Therefore, a suitable network perturbation for changing the signal flow path(s) from an initial state and thereby the absorption probability is to alter the Boolean value taken by one or more of the cross-talk regulation nodes to those specified in Table 3A. We consider two perturbations, *viz*., knocking-off Comp1 – IKK*,^48–50^ and downregulating the inhibitors of apoptosis using smαc – mimetics.^51,52^

After removing the Comp1 – IKK* node along with the interactions it is associated with, we identified the attractors, implemented BM-ProSPR on the perturbed network (ΔComp1 – IKK*) and computed the absorption and steady-state probabilities (Text S6, Supplementary Information). The steady-state probability of ΔComp1 – IKK* network stimulated with TNFα to settle into pro-survival and apoptotic attractors are 0.66 and 0.34, respectively, which amount to a ~20% change compared to those for an unperturbed network stimulated with TNFα (Fig. 7A; Fig. S5-Panel IV, Supplementary Information.) Venn diagram in Fig. 7E shows how the states are distributed into the two basins of attractions. A shift in the cumulative distribution of 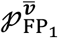 for ΔComp1 – IKK* case towards the ordinate, quantified by the D-statistics^53^ value of ~0.32, indicates that knocking-off Comp1 – IKK* indeed increases the probability of several states in the STG to absorb into apoptosis attractor (Fig. 7F). This demonstrates that knocking-off Comp1 – IKK* can cause a phenotypic switching from prosurvival to apoptosis in an ensemble of cells.

We introduce sma – mimetis into the network by removing the inhibitory action of XIAP causing downregulation of the activation of c3* – p20 by sma c^51,54,55^ and accordingly re-frame the function 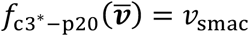. Further, to ensure inhibition of cIAP1/2*,^51^ we reframe the function 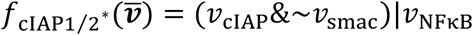. Introduction of smac – mimetics into the network led to a shift in the absorption probability distribution from pro-survival to apoptotic response, indicating phenotype switching in a fraction of cells in an ensemble (Fig. S7-Panel IV, Supplementary Information).

## 3. Discussion and Conclusion

Dynamic cross-talk regulating the TNFR1 signaling mediated pro-survival and apoptotic phenotypic responses to TNFα stimulation is well-known and can be capitalized upon for various therapeutic purposes.^5,56,57^ In this study, using a Boolean dynamic simulation based signal flow path analysis accounting for cell-to-cell variability, we demonstrate that the nested loop involving Comp1 – IKK*, IκB*, IKK*, PI3K, NFκB and Raf1 regulates the cross-talk signaling between TNFR1 network mediated pro-survival and apoptotic responses. We developed and benchmarked a *novel* approach ‘<u>B</u>oolean <u>M</u>odeling based <u>Pr</u>ediction <u>o</u>f <u>S</u>teadystate probability of <u>P</u>henotype <u>R</u>eachability’ (BM-ProSPR) to reliably construct the underlying state transition graph (STG) governing the Boolean dynamics and estimate the absorption probabilities. We accounted for cell-to-cell variability originating due to multitude of signal flow paths using the Random order asynchronous (ROA) Boolean update scheme.

BM-ProSPR employs Temporality^36^ and PageRank^38^ measures to self-learn the extent of evolution of the STG and thereby, aid in identifying the minimum number of permutations required to adequately capture the signal flow paths. For all three considered networks, consisting of small to large number of nodes, BM-ProSPR predicted that a (tiny) fraction of the maximum possible permutations is sufficient for constructing reliable partial STG and computing the absorption probabilities (Fig. 6, Fig. S2, Fig. 7). For example, only 216 out of 17! = 3.55 × 10^14^ permutations are sufficient to reliably compute the absorption and steadystate probabilities of TNFR1 network reaching pro-survival and apoptotic phenotypes when activated with TNFα (Fig. 7). This proves that the proposed algorithm significantly reduces the underlying computational cost. The self-learning nature of BM-ProSPR and the ensuing low computational cost makes it amenable to larger networks for which a complete STG is certainly unavailable. Thus, performing signal flow analysis and thereby distilling out causal structures in large biological networks becomes feasible now.

Our analysis revealed that ~93% of the states reaches apoptosis attractor with a finite absorption probability (Fig. 7C) indicating that a large fraction of the cells in an ensemble has the potential to exhibit TNFR1 signaling mediated apoptotic response. However, the underlying stochasticity in the signal flow path chosen in each of these cells offers only an average steady-state probability of ~21% to reach apoptotic phenotype. Recent experimental studies on primary T-cells,^58^ chondrocytes^59^ and HL60 tissue,^60^ respectively showing 8 – 12% cells committing to apoptosis corroborates our predictions based on the network signal flow analysis. Our analysis proved that perturbing a nested loop by knocking-off Comp1 – IKK^*^could re-wire the signal flow paths to enable phenotype switching from pro-survival to apoptotic response (Fig. 7F). This re-wiring results in a 62% increase in the steady-state probability to reach apoptotic phenotype over that obtained for the unperturbed case. Since Comp1 – IKK* is an important node in TNFR1 signaling,^48,49^ a complete knock-off may not always be desirable. Modulation of the activity of Comp1 – IKK* via inhibition of the Nemo, a subunit of IKK*, has indeed been demonstrated experimentally.^47,61^

Our analysis revealed that after a certain number of permutations, the distribution of PageRank does not change significantly indicating that the topology around some of the states in the STG may have saturated at much lesser number of permutations. Identifying these saturated states while monitoring the STG evolution, for example, using IsoRank,^62^ could help further curtail the computational cost in assembling the state transition matrix. BM-ProSPR algorithm starts with a null STG having all states isolated. Therefore, implementation of BM-ProSPR on very large networks would strongly depend upon the ability to enumerate all possible states and to estimate the absorption probabilities, which can pose a computational challenge.

## 4. Methods

### 4.1 One-step state transition using Random Order Asynchronous update scheme

Consider a state 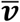. First, a unique permutation sequence is chosen uniformly randomly. Let the first node in the permutation be node *i*. An intermediate state 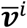 is arrived at by updating the Boolean value of node *i* in state 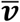 with that resulting from the evaluation of the corresponding Boolean function 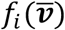. Consider the next node *j* in the permutation sequence. The second intermediate state, say 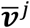, is computed by updating the Boolean value of node *j* in 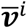 after evaluating the Boolean function 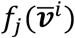. Similarly, intermediate states are generated repeatedly until all nodes in the permutation sequence are exhausted. The final state thus achieved is the one obtained due to a one-step state transition using random order asynchronous update scheme.

### 4.2 Partial logical steady-state analysis (pLSSA)

The network was converted into a list of interactions with the associated logic embedded. This list was parsed into CellNetAnalyzer (CNA).^32^ After fixing Housekeeping nodes with a value of ‘1’, the value corresponding to the relevant input node was specified as ‘1’. The option “*compute logical steady-state*” in CNA was used to identify the nodes attaining partial Logical Steady-State and the corresponding Boolean value.

### 4.3 Identification of Fixed points

Consider any one permutation *q*_1_ out of the *N*! possibilities. Starting from each of the 2*^N^* states ∈ 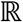, perform one ROA update using *q*_1_ to arrive at those many one-step transitions. The states whose one-step transition leads to the same state, that is, self-loop, are the fixed point attractor. An illustration for identification of fixed points is in Text S2, Supplementary Information.

### 4.4 PageRank

For a certain permutation *q*, using state transition matrix *M^q^*, the PageRank vector was estimated by solving 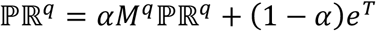 where, *α* = 0.85, *e^T^* is the (column) vector of ones. The PageRank 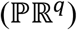 for every permutation was computed using inner-outer iterative scheme.^38^

### 4.5 Kendall-Tau rank correlation

For a certain permutation *q*, the elements of 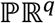 are re-sorted in the same order of placement of the states as per 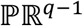, that is, PageRank of the penultimate permutation. Kendall-Tau rank correlation 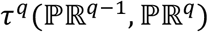 was computed by comparing 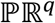 and 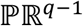 and thereby enumerating the concordant and discordant pairs. A SciPy implementation *scipy.stats.kendalltau*, accessed from Matlab R2018b^®^, was used for this purpose.^63^

### 4.6 Absorption probability

The state transition matrix was re-arranged into a canonical form of 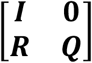 where, *I* is identity matrix that denotes self-loops in FPs, ***R*** is the matrix whose non-zero terms denote the states directly connected to FPs via a one-step state transition, **0** is the zero matrix, and **Q** is the matrix whose non-zero elements show the connection between transient states. Absorption probability 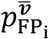 is given by the element corresponding to the state 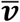 in the i^th^ column of the matrix (***I – Q***)^-1^***R***, where ***I*** is the identity matrix whose dimensions are same as that of ***Q***.^64^The matrix (***I – Q***)^-1^***R*** was calculated using bi-conjugate gradient stabilized method (*bicgstab*) in Matlab R2018b^®^. The *bicgstab* converged usually within 5 to 7 iterations.

## Acknowledgements

This study was funded by the grants CRG/2020/002672 (GV) and MTR/2020/000589 (GV) from Science and Engineering Research Board, Department of Science and Technology, Government of India. We gratefully acknowledge an access to the C-DAC Supercomputing Facility. SS is funded by Department of Biotechnology Junior Research Fellowship (DBT/2017/IIT-B/852).

## List of Supplementary Text

**Text S1:** TNFR1 Signaling network and its Boolean representation

**Text S2:** Identification of fixed point attractors

**Text S3:** Total number of concordant and discordant pairs

**Text S4:** Basin of attraction of the fixed point attractors of the T-LGL apoptosis network

**Text S5:** BM-ProSPR implementation on an 8-node developmental transcription factor network

**Text S6:** Minimum number of permutations for computing state transition matrix for different conditions.

## List of Supplementary Figures

**Figure S1:** Fixed point attractors. (A) One-step state transition between transient states, transient states to FP attractor, FP attractor to FP attractor from a randomly chosen permutations which is mentioned in blue color. The fixed points of the TNFR1 signaling network (Fig. 1, main text) under no stimulation, TNFα stimulation and FasL stimulation are in (B), (C) and (D), respectively.

**Figure S2:** 8-node developmental transcription factor permitting multiple phenotypes. (A) 8-node network having nodes as transcription factors with inhibitory interactions between them. Dependence of (B) Temporality measure 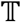, and (C) Fraction of discordant pairs 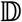 on the permutations. (D) PageRank distribution for *q_l_* = 167 permutation. (E) Distribution of the absorption probability for reaching *p*0 phenotype.

**Figure S3:** FP reachability. (A) Signaling flow path to reach FP. Various paths are shown from a state 01100011 (state ID 117) to reach the fixed point state ID 113. States with yellow color are transient ones and green represents a FP. Transition probability for the one-step state transitions are shown next to the directed interaction in the STG. (B) Reachability to different cell types (phenotypes) from a few states along with the corresponding absorption probabilities. (C) Venn diagram showing sharing of states between the five FPs. The steady-state probability of network’s ability to settle in these five FPs is mentioned next to ellipse.

**Figure S4:** Multiple FP reachability. (A) All signaling flow paths from state 00001110 (state ID 14) that can reach to two different FPs, that is, *p*0 (state 00101110, ID 30) and *p*3 (state 01010000, ID 193). The values next to the arrows capture the corresponding one-step state transition probability **(B)** Comparison of absorption probabilities computed from reliable partial STG with *q_l_* permutations to those of complete STG.

**Figure S5:** Metrics and pro-survival absorption probabilities for the three input and two perturbation conditions. Effect of permutations on the (Panel I) Temporality measure 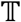 and (Panel II) Fraction of discordant pairs 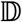. (Panel III) PageRank distribution and (Panel IV) absorption probability distribution for reaching pro-survival phenotype. Note that for the sake of comparison Fig. 7C (main text) is presented as is in Panel IV, TNFα stimulation case.

