## Supplementary Information for "Boolean dynamic modeling of TNFR1 signaling predicts a nested feedback loop regulating the apoptotic response at single-cell level"

Shubhank Sherekar and Ganesh Viswanathan*
Department of Chemical Engineering, Indian Institute of Technology Bombay, Powai Mumbai – 400076

### Table of contents

**S1 Text.** TNFR1 Signaling network and its Boolean representation…………………………[S4]

**S2 Text.** Identification of fixed point attractors……………………………………………..…[S9]

**S3 Text.** Total number of concordant and discordant pairs………………………………[S11]

**S4 Text.** Basin of attraction of the fixed-point attractors of T-LGL apoptosis network……………………………………………………………………………………..[S12]

**S5 Text.** BM-ProSPR implementation on an 8-node developmental transcription factor network……………………………………………………..…………………………[S13]

**S6 Text.** Minimum number of permutations for computing state transition matrix for different conditions……………………………………………………..…………………………[S17]

List of figures

**S1 Fig**: Fixed point attractors…………………………………………………………….…[S10]

**S2 Fig**: 8-node developmental transcription factor permitting multiple phenotypes…………[S13]

**S3 Fig**: FP reachability………..……………………………………………………………[S15]

**S4 Fig**: Multiple FP reachability………..…………………………………………………..…[S16]

**S5 Fig**: Metrics and pro-survival absorption probabilities for the three input and two perturbation conditions. ………..…………………………………………………………………..…[S17]

List of Tables

**S1 Table**: Entities in TNFR1 network (Fig. 1) along with its classification and position in the Boolean state. ………..……………...…………………………………………………..…...[S5]

**S2 Table**: Boolean functions for entities. &, | and ~ operators respectively capture $AND$, $OR$ and $NOT$ operations. ………..……………...…………………………………………………..…...[S6]

**S3 Table**: partial logical steady-state (pLSS) computed for different input conditions in TNFR1 signaling network using CellNetAnalyzer.[29] X means the state of node is "undetermined" from pLSS analysis and can either be 0 or 1……………………………………………………[S7]

**S4 Table**: List of states, sorted in the increasing order of state ID, corresponding to the T-LGL network (Fig. 3) along with absorption probability $\wp^{\overline{v}}_{\mathrm{FP}_1}$. The probability of a state reaching $\mathrm{FP}_2$ $\wp^{\overline{v}}_{\mathrm{FP}_2} = 1 - \wp^{\overline{v}}_{\mathrm{FP}_1}$. Note that the order in which the states are presented in merely for convenience purposes and do not reflect any specific preference to a state or otherwise. …………………[S12]

**S5 Table**: States corresponding to different phenotypic responses…………………………..…[S14]

**S6 Table**: BM-ProSPR predicted minimum number of permutations $q_l$ for reliable estimation of the partial state transition matrix for different input and perturbation conditions on the TNFR1 network………..……………...…………………………………………………..………...[S18]

**S7 Table**: FPs reached by activated TNFR1 signaling network after introduction of perturbations. $FP_1$ and $FP_2$, respectively refers to pro-survival and apoptotic phenotypes. Procedure described in Text S2.1 was used finding the FPs. ……………………………………………..………..[S18]

### Text S1: TNFR1 Signaling network and its Boolean representation

**S1.1**: TNFR1 signaling network

Signaling downstream Tumor Necrosis Factor Receptor 1 (TNFR1) is triggered by the binding of 17 kDa TNFα to it.[1] Following activation, TNFR1 signals both cell death and cell survival via $\mathrm{Comp1}$, a complex consisting of $\mathrm{TRADD}$, $\mathrm{TRAF2}$, $\mathrm{RIP1}$, $\mathrm{cIAP1/2}^*$.[2,3] In the cell-survival arm, the interaction between $\mathrm{IKK}^*$ and $\mathrm{Comp1}$ forms a intermediate complex $\mathrm{Comp1}-\mathrm{IKK}^*$. $\mathrm{Comp1}-\mathrm{IKK}^*$ essentially captures LUBAC complex regulating the $\mathrm{IKK}^*$ activity from $\mathrm{Comp1}$ through the regulatory unit Nemo.[4-6] Activated IκB ($\mathrm{I\kappa B}^*$) is bound to NFκB as $\mathrm{NF\kappa B}-\mathrm{I\kappa B}^*$ in resting cells.[7] Deactivation of $\mathrm{I\kappa B}^*$ by $\mathrm{IKK}^*$ or $\mathrm{Comp1}-\mathrm{IKK}^*$ leads to its disassociation from the $\mathrm{NF\kappa B}-\mathrm{I\kappa B}^*$ complex and thereby releasing the NFκB (functional form).[3] This implies that $\mathrm{I\kappa B}^*$, when present, sequesters the available NFκB and thereby, playing an inhibitory role on NFκB. NFκB up-regulates PI3K,[8,9] which subsequently activates Raf1.[10] Both PI3K and Raf1 promote the transition from the inactive to active forms of IKK,[11] and resulting in a feedback back loop involving these along with $\mathrm{IkB}^*$, NFκB. Downstream, NFκB, being a master regulator, activates various apoptosis inhibitors ($\mathrm{cIAP1/2}$, XIAP, $\mathrm{BCL}-\mathrm{xL}$) and thereby, plays a deciding role in apoptotic response.[7,12] PI3K activates PKB which eventually controls the intrinsic apoptotic response.[13]

In the apoptotic arm of the TNFR1 signalling, $\mathrm{Comp1}$ activates caspase 8 ($\mathrm{c8}^*$) via a series of interactions ($\mathrm{Comp2}$, FADD, c8).[14,15] $\mathrm{c8}^*$ activates the subunit of caspase3 ($\mathrm{c3}^*-\mathrm{p20}$).[16] This activation step is regulated by XIAP via an inhibitory action. While $\mathrm{c3}^*-\mathrm{p20}$ or $\mathrm{c8}^*$ can activate $\mathrm{c3}^*-\mathrm{p17}$, XIAP is known to inhibit both these interactions.[17,18] On the other hand, $\mathrm{c3}^*-\mathrm{p20}$ is also activated via the intrinsic apoptotic pathway originating from Bax, the levels of which is controlled by an inhibitory action exhibited by $\mathrm{BCL}-\mathrm{xL}$ and PI3K.[19-21] Bax activates smac which requires feedback support from $\mathrm{c3}^*-\mathrm{p17}$ to activate $\mathrm{c3}^*-\mathrm{p20}$ in the absence of XIAP.[22-25] $\mathrm{c3}^*-\mathrm{p17}$ executes Apoptosis process by inhibiting downstream protein $\mathrm{Parp}^*$ that inhibits $\mathrm{CAD}^*$ (which is responsible for DNA fragmentation and chromatin condensation).[26]

Fas receptor in presence of FasL leads to apoptotic response by interacting with FADD and forms DISC complex. DISC activates caspase 8 ($\mathrm{c8}^*$) directly through complex formation.[27] All other interactions downstream of $c8^*$ is same as that in TNFR1 signaling.[28] Since most of apoptosis regulating molecular players are shared between Fas and TNFR1 signaling networks, Fas signalling may be considered a positive control for TNFα mediated apoptotic phenotypic response.

**S1.2:** Nodes in the network, Boolean functions and partial Logical steady state analysis

List of included nodes in the TNFR1 network along with the classification, i.e., housekeeping/ input/signaling/output entity are in Table S1. Besides, the order in which the entities are presented in Table S1 specifies the location in the network state ($\bar{\boldsymbol{v}}$) containing the corresponding Boolean value.

The Boolean functions corresponding to the signaling and output entities are in Table S2. Note that while the Boolean values of housekeeping entities are fixed at a certain pre-decided value, those of input nodes are specified *a priori* depending on the stimulation condition considered.

**Table S1:** Entities in TNFR1 network (Fig. 1) along with its classification and position in the Boolean state.

| Name of the entity | Short name | Alignment | Classification |
|---|---|---|---|
| TRADD | TRADD | 1 | Housekeeping |
| RIPK1 | RIP | 2 | Housekeeping |
| TRAF2 | TRAF2 | 3 | Housekeeping |
| FADD | FADD | 4 | Housekeeping |
| $p-14-3-3$ | $p-14-3-3$ | 5 | Housekeeping |
| [c8] | c8 | 6 | Housekeeping |
| [IKK] | IKK | 7 | Housekeeping |
| [cIAP1/2] | cIAP1/2 | 8 | Housekeeping |
| [PARP] | PARP | 9 | Housekeeping |
| TNFα | TNFα | 10 | Input |
| FasL | FasL | 11 | Input |
| TNFR1 | TNFR1 | 12 | Signaling |
| complex 1 | Comp1 | 13 | Signaling |
| complex 2 | Comp2 | 14 | Signaling |
| $c8^{*}-$ complex 2 | $c8^{*}-$ Comp2 | 15 | Signaling |
| Fas | Fas | 16 | Signaling |
| DISC | DISC | 17 | Signaling |
| $c8^{*}-$ DISC | $c8^{*}-$ DISC | 18 | Signaling |
| $c8^{*}$ | $c8^{*}$ | 19 | Signaling |
| cIAP1/2 | $cIAP1/2^{*}$ | 20 | Signaling |
| $c3^{*}-p20$ | $c3^{*}-p20$ | 21 | Signaling |
| $c3^{*}-p17$ | $c3^{*}-p17$ | 22 | Signaling |
| PARP | $PARP^{*}$ | 23 | Signaling |
| CAD | CAD | 24 | Signaling |

| | | | |
|---|---|---|---|
| PI3K | PI3K | 25 | Signaling |
| PKB | PKB | 26 | Signaling |
| Raf1 | Raf1 | 27 | Signaling |
| Bad − 14 − 3 − 3 | Bad − 14 − 3 − 3 | 28 | Signaling |
| BCL − xL | BCL − xL | 29 | Signaling |
| Bax | Bax | 30 | Signaling |
| Smac | smac | 31 | Signaling |
| IκBα | IκB* | 32 | Signaling |
| IKK | IKK* | 33 | Signaling |
| complex 1 − IKK | Comp1 − IKK* | 34 | Signaling |
| FLIP | FLIP | 35 | Signaling |
| XIAP | XIAP | 36 | Signaling |
| NFκB | NFκB | 37 | Output |
| Apoptosis | Apoptosis | 38 | Output |

**Table S2:** Boolean functions for entities. &, | and ~ operators respectively capture $AND$, $OR$ and $NOT$ operations.

| Node Name | Boolean function |
|---|---|
| TNFR1 | TNFα |
| Comp1 | TRADD & TRAF2 & RIP & TNFR1 & cIAP1/2* |
| Comp2 | Comp1 & FADD |
| c8* − Comp2 | c8 & Comp2 |
| Fas | FasL |
| DISC | Fas & FADD |
| c8* − DISC | c8 & DISC |
| c8* | c8* − Comp2 \| (c8* − DISC & ~FLIP) |
| cIAP1/2* | cIAP1/2 \| NFκB \| ~smac |
| c3* − p20 | (smac & c3* − p17 & ~XIAP) \| (c8* & ~XIAP) |
| c3* − p17 | (c3* − p20 & ~XIAP) \| (c8* & ~XIAP) |
| PARP* | PARP & ~c3* − p17 |
| CAD | ~PARP* |
| PI3K | NFκB |
| PKB | PI3K |
| Raf1 | PI3K |
| Bad − 14 − 3 − 3 | P − 14 − 3 − 3 & PKB |
| BCL − xL | NFκB & ~c3* − p17 |
| Bax | ~BCL − xL & ~Bad − 14 − 3 − 3 |
| smac | Bax |
| IκB* | ~IKK* & ~Comp1 − IKK* |
| IKK* | IKK & PI3K & Raf1 |
| Comp1 − IKK* | Comp1 & IKK* |
| FLIP | NFκB |

| | |
|---|---|
| XIAP | NFκB |
| NFκB | ~IκB* |
| Apoptosis | CAD & ~PARP* |

**S1.3:** partial Logical steady-state analysis (pLSSA)

pLSSA was performed on the TNFR1 network for all the cases (including no stimulation) considered using CellNetAnalyzer.[29] Nodes that attained a pLSS for different conditions considered in this study are in Table S3.

**Table S3-** partial logical steady-state (pLSS) computed for different input conditions in TNFR1 signaling network using CellNetAnalyzer.[29] X means the state of node is "undetermined" from pLSS analysis and can either be 0 or 1

| Node ID | Signaling/ Output nodes | pLSS | | | | |
|---|---|---|---|---|---|---|
| | | Basal | TNFα | FasL | TNFα + ΔComp1 − IKK * | TNFα + smac − mimetics |
| 12 | TNFR1 | 0 | 1 | 0 | 1 | 1 |
| 13 | Comp1 | 0 | 1 | 0 | 1 | 0 |
| 14 | Comp2 | 0 | 1 | 0 | 1 | 0 |
| 15 | c8* − Comp2 | 0 | 1 | 0 | 1 | 0 |
| 16 | Fas | 0 | 0 | 1 | 0 | 0 |
| 17 | DISC | 0 | 0 | 1 | 0 | 0 |
| 18 | c8* − DISC | 0 | 0 | 1 | 0 | 0 |
| 19 | c8* | 0 | 1 | X | 1 | 0 |
| 20 | cIAP1/2* | 1 | 1 | 1 | 1 | 0 |
| 21 | c3* − p20 | X | 1 | X | 1 | 1 |
| 22 | c3* − p17 | X | X | X | X | X |
| 23 | PARP* | X | X | X | X | X |
| 24 | CAD | X | X | X | X | X |
| 25 | PI3K | X | X | X | X | X |
| 26 | PKB | X | X | X | X | X |
| 27 | Raf1 | X | X | X | X | X |
| 28 | Bad − 14 − 3 − 3 | X | X | X | X | X |
| 29 | BCL − xL | X | X | X | X | X |
| 30 | Bax | X | X | X | X | X |
| 31 | smac | X | X | X | X | 1 |
| 32 | IkB* | X | X | X | X | X |
| 33 | IKK* | X | X | X | X | X |
| 34 | Comp1 − IKK* | 0 | X | 0 | 0 | 0 |
| 35 | FLIP | X | X | X | X | X |

| | | | | | | |
|---|---|---|---|---|---|---|
| 36 | XIAP | X | X | X | X | X |
| 37 | NFκB | X | X | X | X | X |
| 38 | Apoptosis | X | X | X | X | X |

### Text S2: Identification of fixed point attractors

S2.1: Procedure for identification of fixed points

In order to identify the fixed-point attractors (or fixed points (FP)) in STG, we chose a permutation uniformly randomly and perform a one-step state transition using ROA update (Methods, Main text) of all states in state-space ($\mathbb{R}$). For this purpose, we chose the case of TNFα stimulation. (Without loss of generality, the approach described here is applicable to any stimulation/condition of the network.) In Fig. S1, using the permutation sequence 25,33,10,37,38,27,24,16,32,13,29,20,15,30,22,19,28,35,23,14,21,26,34,17,36,18,31 we show a few one-step transitions. Starting from a state, one-step transition using any permutation can lead to three outcomes, *viz.*,

(i) reach a new state which is not a fixed point. For example, the one-step state transition $\bar{\boldsymbol{v}}_1 \rightarrow \bar{\boldsymbol{v}}_2$ in Fig. S1.

(ii) reach a new state which is a fixed point. For example, the one-step state transition $\bar{\boldsymbol{v}}_2 \rightarrow \bar{\boldsymbol{v}}_3$ in Fig. S1.

(iii) reach the same as the start state. For example, the one-step state transition $\bar{\boldsymbol{v}}_3 \rightarrow \bar{\boldsymbol{v}}_3$, $\bar{\boldsymbol{v}}_4 \rightarrow \bar{\boldsymbol{v}}_4$ in Fig. S1.

One-step transition from a state can lead to itself if and only if the state itself is a FP attractor (case (iii) above). Thus, $\bar{\boldsymbol{v}}_3$ is a FP attractor. Since the output node corresponding to 'Apoptosis' takes a Boolean value of 1 with the other one being 0, this FP is deemed apoptosis attractor. On the other hand, $\bar{v}_4$ is a pro-survival FP as the Boolean value of 'NFκB' output node only takes the Boolean value of 1. We used this approach to identify the FPs for the case of the network (Fig. 1, main text) stimulated with Fas and also unstimulated. These are presented in Fig. S1 below.

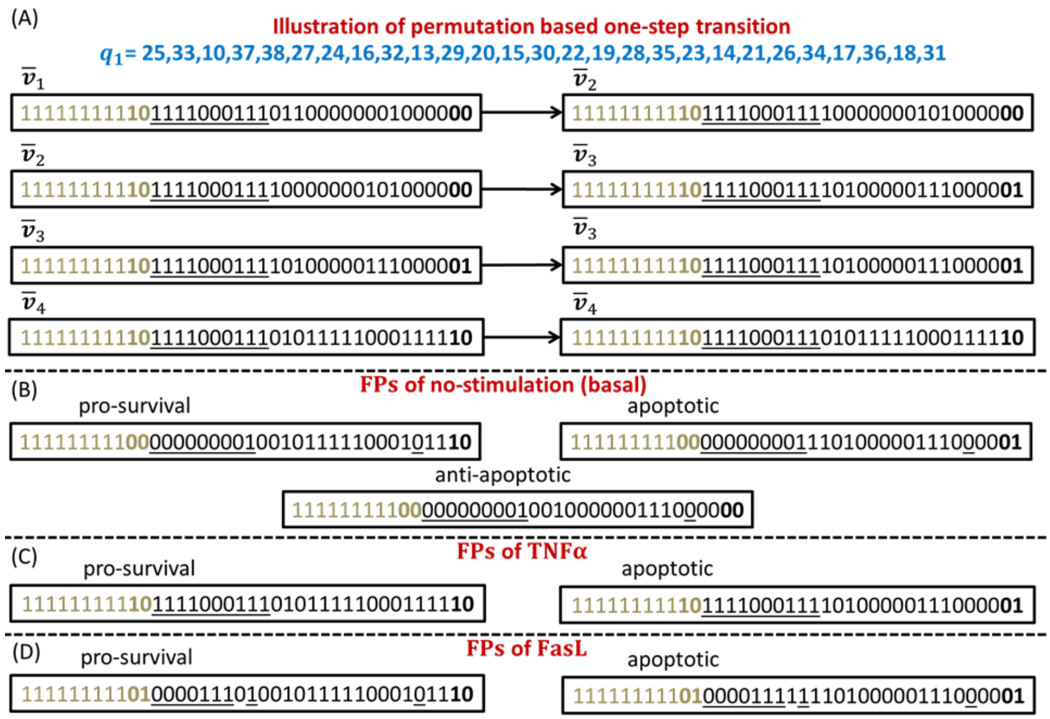

**Figure S1: Fixed point attractors** (A) One-step state transition between transient states, transient states to FP attractor, FP attractor to FP attractor from a randomly chosen permutations which is mentioned in blue color. The fixed points of the TNFR1 signaling network (Fig. 1, main text) under no stimulation, TNFα stimulation and FasL stimulation are in (B), (C) and (D), respectively.

**S2.2**: Permutations for one-step transitions in Fig. 2A

The permutations used for simulating the one-step transitions reported in Fig. 2A are

i) ST1- 14,37,21,28,31,23,34,25,33,18,16,38,17,35,27,29,20,22,26,13,10,30,15,19,32,24,36

ii) ST2- 13,33,34,35,32,29,21,28,20,16,27,10,14,15,24,22,38,26,25,17,23,31,36,37,18,30,19

iii) ST3- 27,26,22,28,37,17,30,36,32,10,24,33,29,35,25,23,16,21,34,20,18,15,38,13,14,31,19

iv) ST4- 37,22,24,20,25,30,23,36,21,15,31,16,28,32,29,34,13,18,38,14,26,19,35,17,33,27,10

v) ST5- 14,21,22,34,28,18,29,16,10,13,36,38,19,24,23,31,30,26,32,15,20,17,27,37,35,33,25

vi) ST6- 25,33,10,37,38,27,24,16,32,13,29,20,15,30,22,19,28,35,23,14,21,26,34,17,36,18,31

vii) ST7- 17,28,18,35,31,26,21,27,32,10,34,13,29,25,19,23,36,16,22,20,37,15,38,24,30,33,14

**Text S3: Total number of concordant and discordant pairs**

Kendall's Tau rank correlation[30] was used as a measure to compare PageRank order of states in STG with subsequent number of permutations. For this purpose, the concordant and discordant pairs based on a comparison of PageRank of states in STG at permutation $q$ with that for $(q-1)^{th}$ permutations are identified. Since the number of states in the STG corresponding to the network having $N$ entities is $2^N$, total number of pairs is $\binom{2^N}{2}$. Therefore, the sum of concordant $(C)$ and discordant $(D)$ pairs, irrespective of the chosen permutation for which the Kendall's Tau rank correlation is estimated, is given by

$$C^q + D^q = \binom{2^N}{2} = \frac{2^N(2^N-1)}{2} = 2^{N-1}(2^N - 1). \quad \text{[S4.1]}$$

**Text S4: Basin of attraction of the fixed point attractors of the T-LGL apoptosis network**

A state can reach either one or both the fixed points $\mathrm{FP}_1$ and $\mathrm{FP}_2$. The probability of reaching an attractor from a certain state is its absorption probability. In Table S4, we show the absorption probability $p^{\bar{v}}_{\mathrm{FP}_1}$ of all states reaching $\mathrm{FP}_1$. Those having an absorption probability of 1 belong exclusively to the basin of attraction of that attractor. For example, states 000001 (state ID 2, Table S4), 001011 (state ID 12, Table S4) belong exclusively to the basin of attraction of $\mathrm{FP}_1$. On the other hand, states 011111, 110000 and 110110 (state IDs 32, 49, and 55, respectively in Table S4) belong exclusively to the basin of attraction of $\mathrm{FP}_2$. All other states belong to both basin of attraction of $\mathrm{FP}_1$ and $\mathrm{FP}_2$ with $0 < p^{\bar{v}}_{\mathrm{FP}_1} = 1 - p^{\bar{v}}_{\mathrm{FP}_1} < 1$.

**Table S4:** List of states, sorted in the increasing order of state ID, corresponding to the T-LGL network (Fig. 3) along with absorption probability $p^{\bar{v}}_{\mathrm{FP}_1}$. The probability of a state reaching $\mathrm{FP}_2$ $p^{\bar{v}}_{\mathrm{FP}_2} = 1 - p^{\bar{v}}_{\mathrm{FP}_1}$. Note that the order in which the states are presented in merely for convenience purposes and do not reflect any specific preference to a state or otherwise.

| ID | State | $p^{\bar{v}}_{\mathrm{FP}_1}$ | ID | State | $p^{\bar{v}}_{\mathrm{FP}_1}$ | ID | State | $p^{\bar{v}}_{\mathrm{FP}_1}$ |
|---|---|---|---|---|---|---|---|---|
| 1 | 000000 | 0.235 | 23 | 010110 | 1 | 45 | 101100 | 1 |
| 2 | 000001 | 1 | 24 | 010111 | 0.907 | 46 | 101101 | 0.649 |
| 3 | 000010 | 1 | 25 | 011000 | 1 | 47 | 101110 | 1 |
| 4 | 000011 | 0.665 | 26 | 011001 | 0.501 | 48 | 101111 | 1 |
| 5 | 000100 | 1 | 27 | 011010 | 1 | 49 | 110000 | 0 |
| 6 | 000101 | 0.725 | 28 | 011011 | 1 | 50 | 110001 | 1 |
| 7 | 000110 | 1 | 29 | 011100 | 1 | 51 | 110010 | 0.934 |
| 8 | 000111 | 0.919 | 30 | 011101 | 1 | 52 | 110011 | 1 |
| 9 | 001000 | 1 | 31 | 011110 | 1 | 53 | 110100 | 0.499 |
| 10 | 001001 | 0.694 | 32 | 011111 | 0 | 54 | 110101 | 1 |
| 11 | 001010 | 1 | 33 | 100000 | 1 | 55 | 110110 | 0 |
| 12 | 001011 | 1 | 34 | 100001 | 0.811 | 56 | 110111 | 1 |
| 13 | 001100 | 1 | 35 | 100010 | 1 | 57 | 111000 | 0.548 |
| 14 | 001101 | 1 | 36 | 100011 | 0.500 | 58 | 111001 | 1 |
| 15 | 001110 | 1 | 37 | 100100 | 1 | 59 | 111010 | 0.859 |
| 16 | 001111 | 0.966 | 38 | 100101 | 0.553 | 60 | 111011 | 1 |
| 17 | 010000 | 1 | 39 | 100110 | 1 | 61 | 111100 | 0.587 |
| 18 | 010001 | 0.166 | 40 | 100111 | 0.863 | 62 | 111101 | 1 |
| 19 | 010010 | 1 | 41 | 101000 | 1 | 63 | 111110 | 0.592 |
| 20 | 010011 | 0.626 | 42 | 101001 | 0.296 | 64 | 111111 | 0.894 |
| 21 | 010100 | 1 | 43 | 101010 | 1 | | | |
| 22 | 010101 | 0.703 | 44 | 101011 | 0.835 | | | |

**Text S5: BM-ProSPR implementation on an 8-node developmental transcription factor network**

In order to demonstrate BM-ProSPR's ability to compute absorption probabilities to reach multiple phenotypes (FPs), we consider an 8-node developmental transcription factor network regulating the spinal cord ventrilization (Fig. S2A).[31] The ventrilization network permits 5 progenitor cell types depending upon the activated or inactivated state of different nodes. We frame Boolean

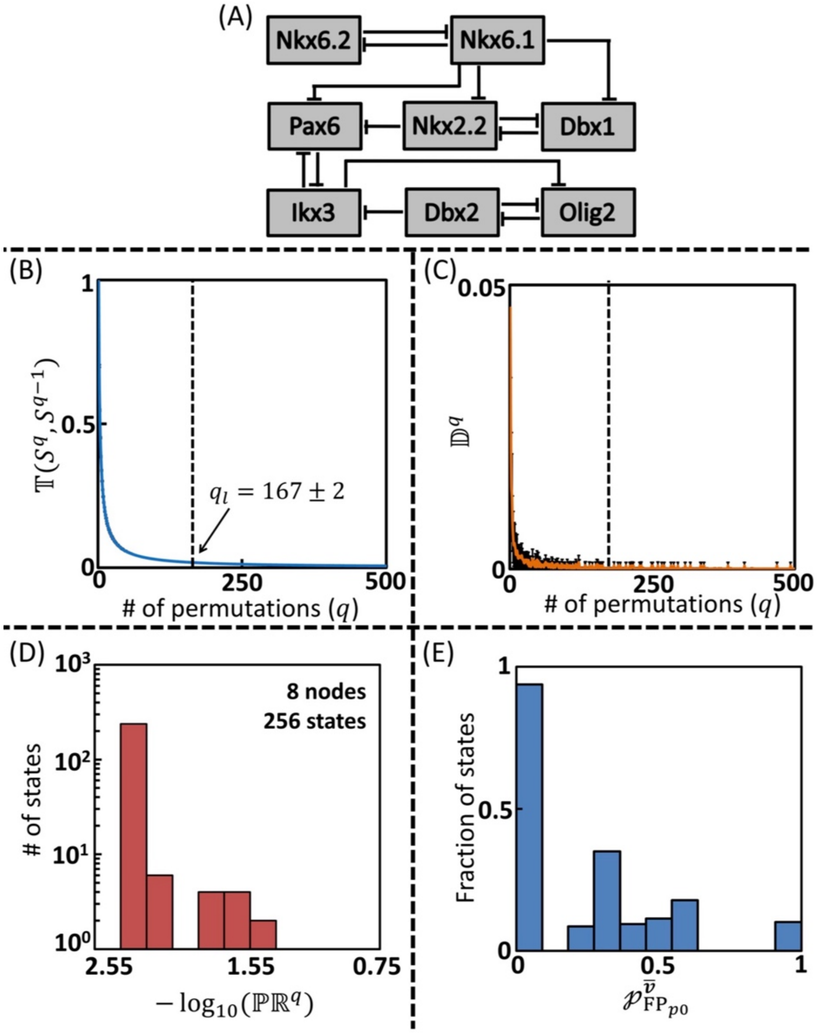


**Figure S2: 8-node developmental transcription factor permitting multiple phenotypes.** (A) 8- node network having nodes as transcription factors with inhibitory interactions between them. Dependence of (B) Temporality measure $\mathbb{T}$, and (C) Fraction of discordant pairs $\mathbb{D}$ on the permutations. (D) PageRank distribution for $q_l = 167$ permutation. (E) Distribution of the absorption probability for reaching $p0$ phenotype.

functions for every node capturing the inhibitory actions on it by other entities in the network. For example, the Boolean function corresponding to Pax6 , inhibited by Nkx6.1, Nkx2.2, Ikx3, is

$$f_{\mathrm{Pax6}}(\overline{\boldsymbol{v}}) = \sim(v_{\mathrm{Nkx6.1}} | v_{\mathrm{Nkx2.2}} | v_{\mathrm{Ikx3}}) \qquad [S5.1]$$

where ~ and | represent NOT and OR logic, respectively. The fixed points (FPs) $p3, pMN, p2, p1$ and $p0$ of the network correspond to the five cell types and the respective states are in Table S5.

**Table S5:** States corresponding to different phenotypic responses.

Dorsal ⟶ Ventral

| TFs↓ Cells→ | $p3$ | $pMN$ | $p2$ | $p1$ | $p0$ |
|---|---|---|---|---|---|
| **Nkx6.2** | 0 | 0 | 0 | 1 | 0 |
| **Nkx6.1** | 1 | 1 | 1 | 0 | 0 |
| **Pax6** | 0 | 1 | 1 | 1 | 1 |
| **Nkx2.2** | 1 | 0 | 0 | 0 | 0 |
| **Dbx1** | 0 | 0 | 0 | 0 | 1 |
| **Ikx3** | 0 | 0 | 1 | 1 | 1 |
| **Dbx2** | 0 | 0 | 0 | 1 | 1 |
| **Olig2** | 0 | 1 | 0 | 0 | 0 |

We implemented the BM-ProSPR algorithm on this network. STG of the network consists of $2^8 = 256$ states and 50 randomized state transition graph constructions were considered. Temporality measure and fraction of pairs having discordant PageRank across successive permutations show a decreasing trend with increase in permutations (Fig. S2B and S2C). PageRank distribution is shown in Fig. S2D.

Only $q_l = 167 \pm 2$ permutations out of $q_{max} = 40320$ were required for reliable estimation of the partial state transition matrix $M^{q_l}$. While the complete STG will contain $40320 \times 256 = 1{,}03{,}21{,}920$ one-step state transitions, only a tiny fraction of $0.4\% = 170 \times 256 = 43520$ is needed for finding $\mathbb{S}^{q_l}$ required to estimate $M^{q_l}$. A histogram of absorption probability of $p0$ phenotype (Fig. S2E) clearly shows that a fraction of states belongs exclusively to the basin of attraction $p0$ and also several having $0 < \wp_{p0}^{\overline{v}} < 1$ indicating that these states could reach multiple phenotypes. In Fig. S3A, as an example, we show all signal paths originating from the state $01100011$ (state ID 117) and terminating only in the FP $pMN$ (state ID 113). While the probability of reaching $pMN$ via signal path $117 \rightarrow 51 \rightarrow 115 \rightarrow 113$ is $0.19 \times 0.52 \times 1.0 = 0.0988$, that for $117 \rightarrow 50 \rightarrow 113$ is $0.21 \times 0.49 = 0.1029$. These probabilities summed over all signal paths led to an absorption probability $\wp_{pMN}^{117}$ of 1, indicating that state 117 belongs exclusively to the basin of attraction of $pMN$. Such sub-graphs of partial STG starting from a state and culminating in different FPs were used to identify the corresponding absorption probabilities. In Fig. S4A, we present an example of state $00001110$ absorbing into FPs $p0$ and $p3$ along with the one-step transition probabilities.

In order to enable placing the states into the basin of attraction of the five FPs (Table S5), using $\mathcal{M}^{q_l}$, we estimated the absorption probability of each the states to absorb into these fixed points (Fig. S3A). While some states such as $00001110$ could absorb into FPs $p0$ and $p3$ with equal probabilities, many others such as state $00000000$ fall into the basin of attraction of all five FPs with a finite probability (Fig. S3B). Thus, all states in the state space of the STG can be classified according to their absorption probability towards the five FPs. Venn diagram (Fig. S3-C) shows the statistics of the states absorbing into different FPs. Based on these absorption probabilities, we estimated the steady-state probability of the network's ability to settle into the five FPs (Fig. S3C). A comparison with those computed using the complete state transition matrix shows that BM-ProSPR algorithm enables accurate estimation of the steady-state probabilities using a significantly smaller number of Boolean dynamics simulations (Fig. S4B). This clearly demonstrates that the proposed algorithm can reliably lead to identifying the minimum number of permutations $q_l$ required to estimate $\mathcal{M}^{q_l}$ in order to find the steady-state probabilities of different FPs.

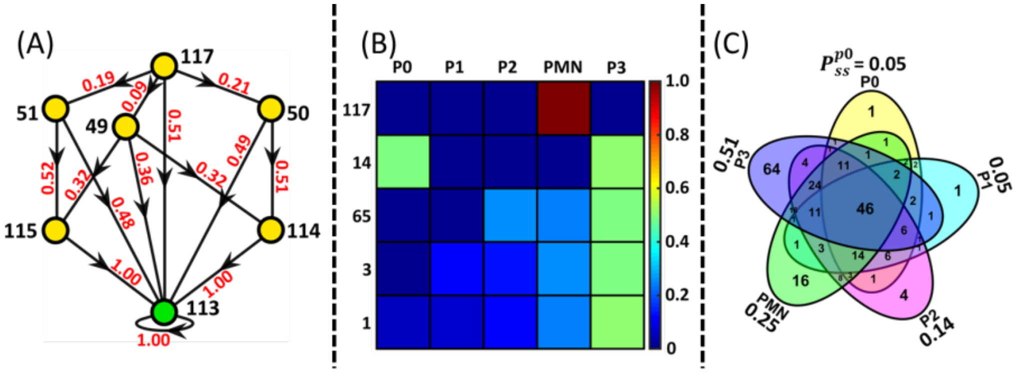


**Figure S3: FP reachability.** (A) Signaling flow path to reach FP: Various paths are shown from a state $01100011$ (state ID 117) to reach the fixed point state ID 113. States with yellow color are transient ones and green represents a FP. Transition probability for the one-step state transitions are shown next to the directed interaction in the STG. (B) Reachability to different cell types (phenotypes) from a few states along with the corresponding absorption probabilities. (C) Venn diagram showing sharing of states between the five FPs. The steady-state probability of network's ability to settle in these five FP is mentioned next to ellipse.

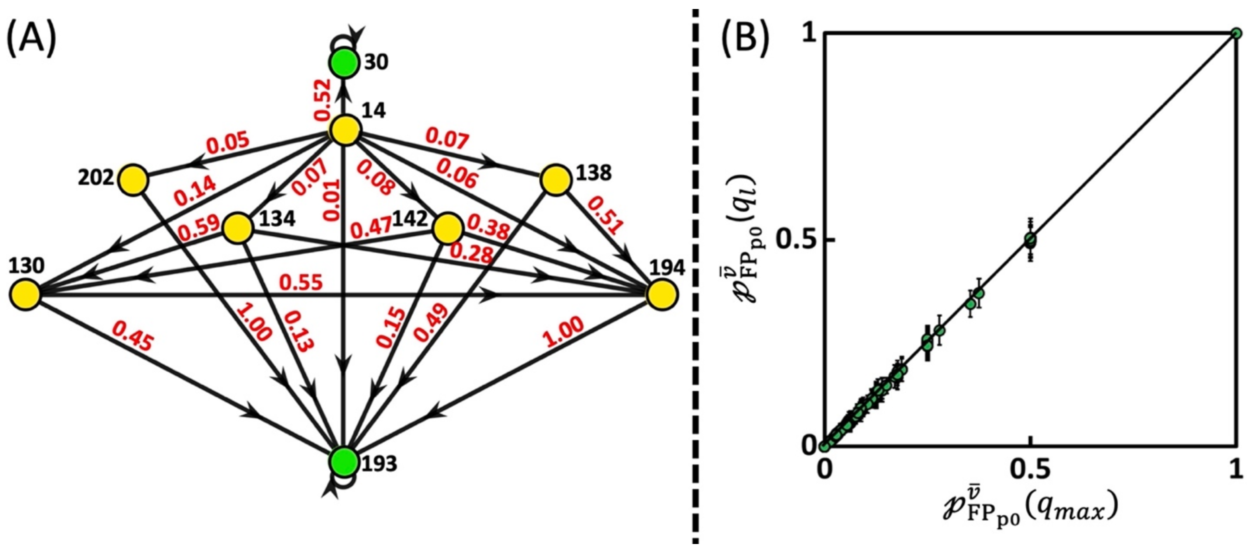


**Figure S4: Multiple FP reachability.** (A) All signaling flow paths from state 00001110 (state ID 14) that can reach to two different FPs, that is, $p0$ (state 00101110, ID 30) and$p3$ (state 01010000, ID 193). The values next to the arrows capture the corresponding one-step state transition probability **(B)** Comparison of absorption probabilities computed from reliable partial STG with $q_l$ permutations to those of complete STG.

**Text S6: Minimum number of permutations for computing state transition matrix for different conditions.**

Temporality measure and the fraction of discordant pairs as a function of the number of permutations for the three input conditions, *viz*., no stimulation (basal), TNFα stimulation, FasL stimulation are in the first three columns of Fig. S5. The minimum number of permutations required to reliably estimate the state transition matrix corresponding to the partial STG of the TNFR1 network Boolean model (Fig. 1) for these cases are in Table S6.

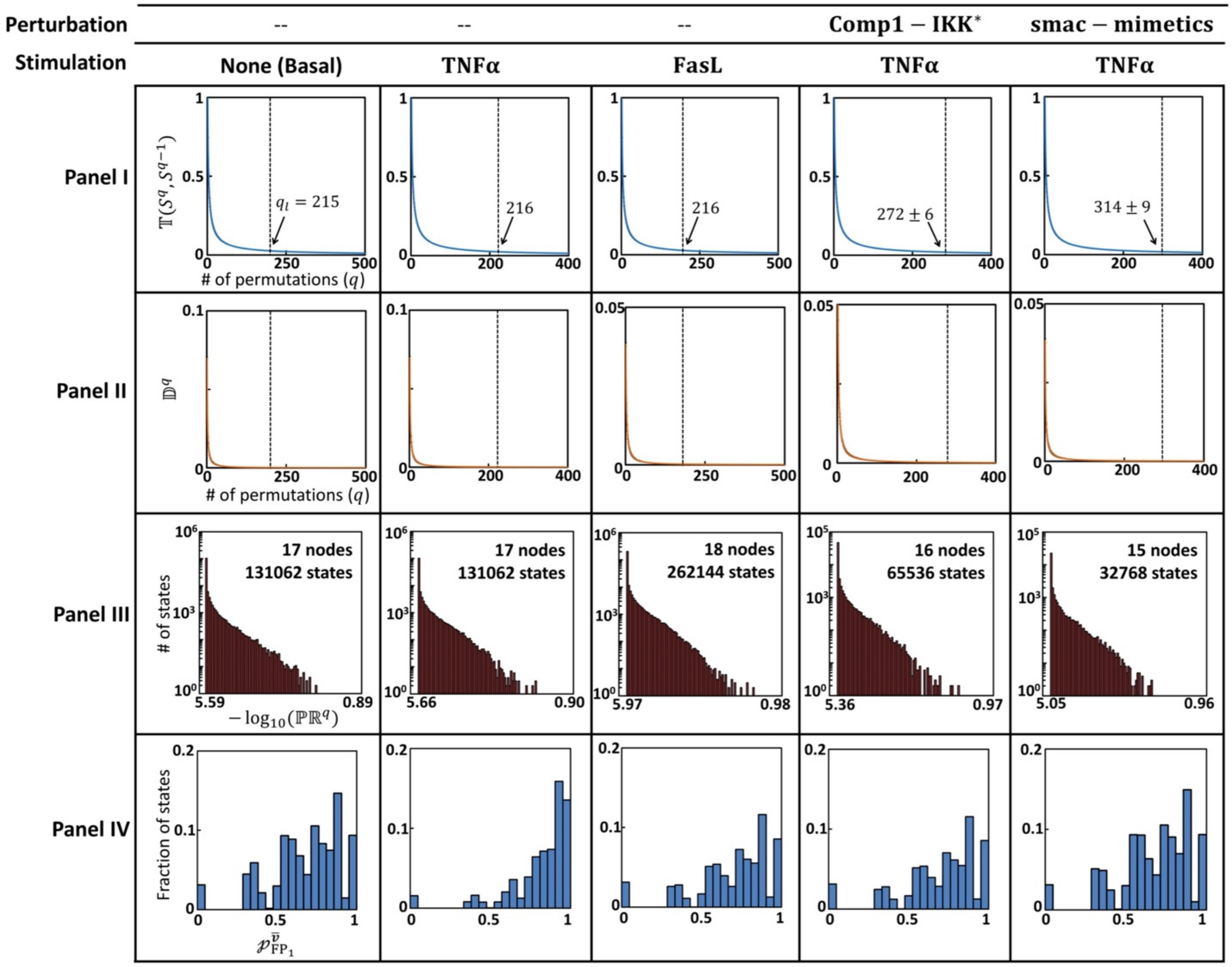


**Figure S5: Metrics and pro-survival absorption probabilities for the three input and two perturbation conditions.** Effect of permutations on the (Panel I) Temporality measure $\mathbb{T}$ and (Panel II) Fraction of discordant pairs $\mathbb{D}$. (Panel III) PageRank distribution and (Panel IV) absorption probability distribution for reaching pro-survival phenotype. Note that for the sake of comparison Fig. 7C (main text) is presented as is in Panel IV, TNFα stimulation case.

For the case of two perturbed networks, *viz*., $\Delta\text{Comp1}-\text{IKK}^*$ and $\text{smac}-\text{mimetics}$, stimulated with TNFα, we first identified the attractors using the procedure in Text S2.1. The attractors of these perturbed networks are in Table S7. Dependence of the temporality measure and the fraction of discordant pairs on $q$ are in Fig. S5, columns 4 and 5. Note that knocking off of $\text{Comp1}-\text{IKK}^*$ does not alter the basal response as Comp1 always takes a Boolean value of 0 in the absence of stimulation.

**Table S6**: BM-ProSPR predicted minimum number of permutations $q_l$ for reliable estimation of the partial state transition matrix for different input and perturbation conditions on the TNFR1 network.

| | | $q_l$ |
|---|---|---|
| Input | Basal | 215 |
| | TNF$\alpha$ | 216 |
| | FasL | 216 |
| TNF$\alpha$ +perturbation | $\Delta Comp1-IKK^*$ | 272±6 |
| | $Smac\ mimetics$ | 314±9 |

**Table S7**: FPs reached by activated TNFR1 signaling network after introduction of perturbations. $\text{FP}_1$ and $\text{FP}_2$, respectively refers to pro-survival and apoptotic phenotypes. Procedure described in Text S2.1 was used finding the FPs.

| ID | Entity | $\Delta\text{Comp1}-\text{IKK}^*$ | | $\text{smac}-\text{mimetics}$ | |
|---|---|---|---|---|---|
| | | $\bar{v}_{\text{FP}_1}$ | $\bar{v}_{\text{FP}_2}$ | $\bar{v}_{\text{FP}_1}$ | $\bar{v}_{FP_2}$ |
| 1 | TRADD | 1 | 1 | 1 | 1 |
| 2 | RIP | 1 | 1 | 1 | 1 |
| 3 | TRAF2 | 1 | 1 | 1 | 1 |
| 4 | FADD | 1 | 1 | 1 | 1 |
| 5 | $\text{p}-14-3-3$ | 1 | 1 | 1 | 1 |
| 6 | c8 | 1 | 1 | 1 | 1 |
| 7 | IKK | 1 | 1 | 1 | 1 |
| 8 | cIAP1/2 | 1 | 1 | 1 | 1 |
| 9 | PARP | 1 | 1 | 1 | 1 |
| 10 | TNFα | 1 | 1 | 1 | 1 |
| 11 | FasL | 0 | 0 | 0 | 0 |
| 12 | TNFR1 | 1 | 1 | 1 | 1 |
| 13 | Comp1 | 1 | 1 | 0 | 0 |
| 14 | Comp2 | 1 | 1 | 0 | 0 |
| 15 | $\text{c8}^*-\text{Comp2}$ | 1 | 1 | 0 | 0 |

| | | | | | |
|---|---|---|---|---|---|
| 16 | Fas | 0 | 0 | 0 | 0 |
| 17 | DISC | 0 | 0 | 0 | 0 |
| 18 | $c8^{*} - DISC$ | 0 | 0 | 0 | 0 |
| 19 | $c8^{*}$ | 1 | 1 | 0 | 0 |
| 20 | $cIAP1/2^{*}$ | 1 | 1 | 0 | 0 |
| 21 | $c3^{*} - p20$ | 1 | 1 | 1 | 1 |
| 22 | $c3^{*} - p17$ | 0 | 1 | 0 | 1 |
| 23 | $PARP^{*}$ | 1 | 0 | 1 | 0 |
| 24 | CAD | 0 | 1 | 0 | 1 |
| 25 | PI3K | 1 | 0 | 1 | 0 |
| 26 | PKB | 1 | 0 | 1 | 0 |
| 27 | Raf1 | 1 | 0 | 1 | 0 |
| 28 | $Bad - 14 - 3 - 3$ | 1 | 0 | 1 | 0 |
| 29 | $BCL - xL$ | 1 | 0 | 1 | 0 |
| 30 | Bax | 0 | 1 | 0 | 1 |
| 31 | smac | 0 | 1 | 1 | 1 |
| 32 | $I\kappa B^{*}$ | 0 | 1 | 0 | 1 |
| 33 | $IKK^{*}$ | 1 | 0 | 1 | 0 |
| 34 | $Comp1 - IKK^{*}$ | 0 | 0 | 0 | 0 |
| 35 | FLIP | 1 | 0 | 1 | 0 |
| 36 | XIAP | 1 | 0 | 1 | 0 |
| 37 | $NF\kappa B$ | 1 | 0 | 1 | 0 |
| 38 | Apoptosis | 0 | 1 | 0 | 1 |
